# Pharmacological rescue of cilia trafficking defects in IFT140 retinal organoid and RPE models of retinal dystrophy

**DOI:** 10.64898/2026.04.29.720656

**Authors:** Julio C. Corral-Serrano, Yi Jiang, Nele Schwarz, Sara E. Nieuwenhuis, Kelly Ziaka, Siobhan Guilfoyle, Rosellina Guarascio, Adriana Bakoulina, Marian Seda, Jeshmi Jeyabalan Srikaran, Daniele Ottaviani, Esben Lorentzen, Isabelle Perrault, Alison J. Hardcastle, Tina Beyer, Dagan Jenkins, Michael E. Cheetham

## Abstract

Pathogenic variants in *IFT140* are associated with a spectrum of syndromic and non-syndromic ciliopathies, with retinal degeneration as a common feature. Despite advances in understanding IFT140 function across various tissues, human retina-specific models are lacking. Here, we show that knock-in mice homozygous for the *IFT140* patient variant c.932A>G (p.Y311C) did not develop retinal degeneration, while mice with the homozygous variant c.1451C>T (p.T484M), associated with non-syndromic retinal dystrophy, were embryonic lethal. Therefore, to understand the effect of these variants on retinal homeostasis, we generated novel human *in vitro* models of IFT140-associated retinal dystrophy, including CRISPR/Cas9 IFT140 knock-out (*IFT140^KO^*) induced pluripotent stem cells (iPSC) and patient-derived iPSC retinal pigment epithelium (iPSC-RPE) and retinal organoids (iPSC-ROs). *IFT140^KO^* iPSC-RPE cells display stubby cilia compared to isogenic controls, while *IFT140^T484M/T484M^*patient-derived iPSC-RPE cells exhibit slightly shorter cilia and cilia tip protein accumulation. Both *IFT140^KO^* and *IFT140^T484M/T484M^* iPSC-ROs show accumulation of cilia proteins at the connecting cilium and outer segment of photoreceptors, and mislocalization of rhodopsin to the inner segments and outer nuclear layer. Pharmacological screening of compounds previously reported to improve cilia structure identified the flavonoid eupatilin as the most effective molecule. Treatment with eupatilin improved cilium length and IFT traffic in iPSC-RPE, and IFT traffic and rhodopsin localization in iPSC-ROs. These findings emphasize the importance of human stem cell derived models to investigate tissue specific disease mechanisms and highlight the therapeutic potential of eupatilin to ameliorate cilia defects in retinal tissue.

## Introduction

Primary cilia are microtubule-based organelles found in most cells that function as a sensor of the extracellular environment and are important in many signaling pathways (Mill et al., 2023). Pathogenic variants in cilia genes cause a range of diseases called ciliopathies, that affect different organs, with retinal degeneration as a common feature. Importantly, around 20% of inherited retinal diseases are associated with cilia genes, which are termed retinal ciliopathies (Bujakowska et al., 2017; Rivolta et al., 2025). The outer segments (OS) of photoreceptor cells are modified cilia arranged as a stack of membranous discs that act as light-sensing organelles. A short ciliary bridge, called the connecting cilium, connects the OS to the photoreceptor inner segment (IS), where OS proteins are synthesized. Intraflagellar transport (IFT) along the connecting cilium is therefore crucial for the maintenance and function of photoreceptor OS (Gupta & Pazour, 2024). Currently, there is a need for more detailed studies on the function of the photoreceptor cilium to improve our understanding of retinal ciliopathy disease mechanisms and to develop effective therapies.

IFT140 is part of the IFT-A complex, a protein complex that mediates cilia entry and retrograde trafficking of membrane proteins from the tip of the cilium to the base (Hirano et al., 2017). Recently, a number of studies have revealed the 3D structure of the IFT-A complex (Hesketh et al., 2022; Jiang et al., 2023; Lacey et al., 2023, 2024; Meleppattu et al., 2022; Nachury, 2022). IFT140 contains two tandem WD40 β-propellers and an array of 20 tetratricopeptide repeats (TPR), which are important for IFT complex interaction as well as cargo binding (Hesketh et al., 2022). *IFT140* variants can be classified based on their effect on cargo delivery or IFT train assembly (Ma et al., 2023), for example, *IFT140* missense variants can affect IFT-A complex integrity (Beyer et al., 2025). Biallelic pathogenic variants in *IFT140* are associated with ciliopathies, including Mainzer-Saldino syndrome (MSS) (Geoffroy et al., 2018; Helm et al., 2017; Perrault et al., 2012; Yeh et al., 2022), Jeune asphyxiating thoracic dystrophy (JATD) (Schmidts et al., 2013), non-syndromic Retinitis Pigmentosa (RP), non-syndromic Leber Congenital Amaurosis (LCA) (Hull et al., 2016; Xu et al., 2015), Sensenbrenner syndrome and cranioectodermal dysplasia (CED) (Bayat et al., 2017; Walczak-Sztulpa et al., 2020, 2022). In contrast, monoallelic pathogenic variants in *IFT140* have been recently described as a frequent cause of autosomal dominant polycystic kidney disease (adPKD) (Dordoni et al., 2024; Fujimaru et al., 2024; Griffiths et al., 2026; Salhi et al., 2024; Seeman et al., 2024; Senum et al., 2022). Together, this broad phenotypic spectrum underscores the pleiotropic and tissue-specific functions of IFT140.

Previous studies across multiple model systems have established IFT140 as a critical regulator of cilia function in diverse tissues. Early work in *Trypanosoma* demonstrated that IFT140 loss results in shortened flagella with accumulation of IFT proteins, consistent with defective retrograde transport (Absalon et al., 2008). Similarly, in mammalian systems, disruption of IFT140 leads to cilia defects characterized by accumulation of IFT components such as IFT88 at the cilia tip (Oud et al., 2018). Mouse models have linked IFT140 dysfunction to skeletal ciliopathies resembling Joubert syndrome (JS) (Miller et al., 2013) and to renal cystic disease (Jonassen et al., 2012), highlighting its broad physiological role and its role in normal embryonic development (Francis et al., 2023). In the mouse retina, conditional models show that IFT140 disruption leads to mislocalisation of opsins to the IS, indicating defective protein trafficking (Crouse et al., 2014).

Despite these advances in understanding IFT140 function, there are no models that investigate the effects of IFT140 variants in human retinal tissue. To address this, we generated and characterized novel iPSC-derived models of IFT140-associated retinal disease. Using these systems, we identify defects in cilia structure and trafficking that can be improved by pharmacological intervention. This work highlights the importance of human stem cell derived models for investigating retinal disease associated with ciliopathy genes.

## Results

### *Ift140* knock-in mice with pathogenic missense variants do not develop overt retinal degeneration

To investigate retinal disease mechanisms associated with *IFT140* variants *in vivo*, we generated heterozygous, compound heterozygous and homozygous knock-in mice with different *Ift140* variants analogous to those identified in patients: c.932A>G (p.Y311C), which is reported in MSS patients as a compound heterozygous allele with the frameshift variant c.857-862delTTGA (p.I286Lfs*6) (Perrault et al., 2012), and the non-syndromic RP variant c.1451C>T (p.T484M) (Hull et al., 2016; Xu et al., 2015) (**Figure 1A and Supplementary Figure 1**). Y311C homozygous mice (*Ift140^Y311C/Y311C^*) were produced at the expected mendelian ratios and appeared healthy with no signs of syndromic disease. Strikingly, no homozygous mice from the T484M model (*Ift140^T484M/T484M^*) were born, suggesting issues with embryonic development. Only heterozygous *Ift140^T484M/+^* or wild-type offspring were produced from 11 litters. Examination of the embryos at embryonic day 11.5 (E11.5) revealed features of a severe ciliopathy including anophthalmia, exencephaly and malformation of the limbs (**Figure 1B**), confirming the embryonic lethality in *Ift140^T484M/T484M^*. Staining for cilia makers in mouse embryonic fibroblasts (MEFs) obtained from these embryos revealed no axoneme formation in the homozygous *Ift140^T484M/T484M^* MEFs, whereas cilia formation in *Ift140^T484M/+^*MEFs was the same as control MEFs (**Figure 1C-E and Supplementary Figure 1**). The *Ift140^T484M/+^* MEFs showed normal distribution of IFT88 along the cilium, which was not possible to assess in the *Ift140^T484M/T484M^* MEFs, due to the absence of the axoneme (**Supplementary Figure 1**). The *Ift140^T484M/+^* mice did not show any changes in ONL thickness at 3 or 5 months of age (**Figure 1D**). Similarly, *Ift140^Y311C/+^* and *Ift140^Y311C/Y311C^* mice did not show any significant changes in ONL thickness at 3, 5 and 7 months (**Figure 1D** and **Supplementary Figure 1**). Compound heterozygous mice (*Ift140^Y311C/T484M^*) were produced to investigate if the combination of alleles would reveal a more severe phenotype; however, they appeared healthy with no signs of syndromic disease. *Ift140^Y311C/T484M^* mice also had no reduction in ONL thickness at 5 or 7 months or at the latest time point measured of 12 months (**Figure 1E** and **Supplementary Figure 1**). None of the Ift140 models (*Ift140^Y311C/+^*, *Ift140^Y311C/Y311C^*and *Ift140^Y311C/T484M^*) analyzed showed a reduction in their retinal function compared to control animals by ERG (a and b wave) at up to 8 months of age (**Supplementary Figure 1**).

**Figure 1.**
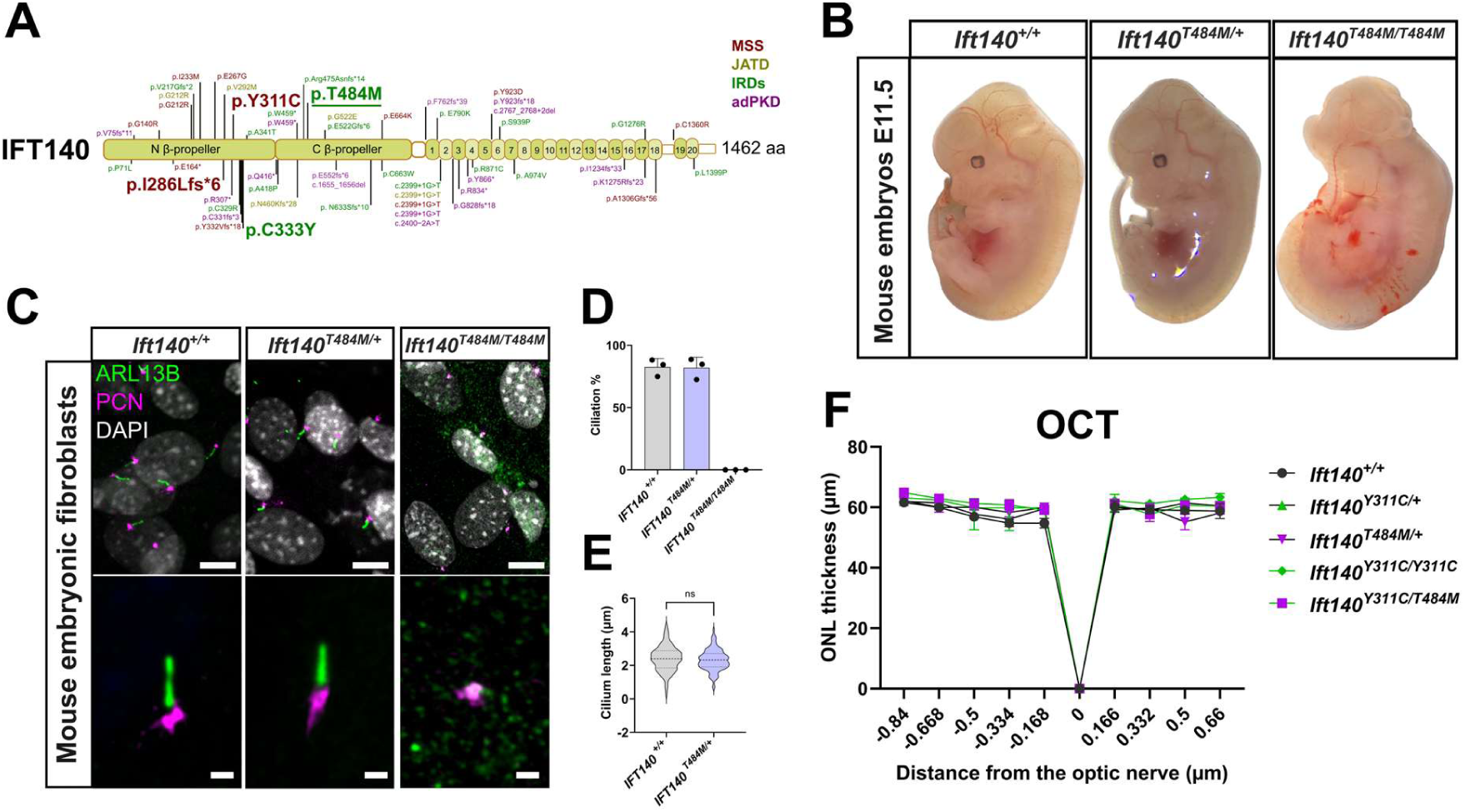
Homozygous *Ift140* T484M knock-in mice have severe cilia defects and are embryonic lethal. **A)** Schematic representation of the IFT140 protein showing all reported variants to date, with the missense variants modelled in this study marked in bold. Disease associations are color coded, MSS (red) JATD, (yellow), IRD (green), aPKD (purple). The T484M variant was also modelled for our human stem cell models studies (underlined). **B)** Images of mouse embryos of each T484M genotype, showing *Ift140^T484M/T484M^* embryos have anophthalmia, exencephaly and malformation of the limbs, whereas *Ift140^T484M/+^* embryos develop normally. **C)** Mouse embryonic fibroblasts were stained with ARL13B (cilia membrane, green) and PCN (basal body, magenta) showing absence of cilia formation in the *Ift140^T484M/T484M^* mice. DAPI (grey) stains the nucleus. Scale bar: 10 µm for the upper images and 1 µm for the bottom images. Two independent mouse embryonic *Ift140^T484M/T484M^* fibroblast lines were analyzed. **D)** Quantification of cilia incidence by measuring ARL13B elaboration from the basal body (n=3, >150 cilia counted). **E)** Cilium length measured for ARL13B from the basal body (*Ift140^+/+^* n=126 cilia; *Ift140^T484M/+^*n=115 cilia). **F)** Optical coherence tomography (OCT) of the outer nuclear layer (ONL) thickness for the different *Ift140* genotypes. The number of mice analyzed for each genotype, at the latest time point analyzed, was as follows: *IFT140^+/+^* (7 months; n=2); *IFT140^Y311C/Y311C^* (7 months; n=3); *IFT140^Y311C/Y311C^* (12 months; n=2); *IFT140^T484M/+^* (5 months; n=4); *IFT140^Y311C/T484M^* (12 months; n=2).

### Cilia retrograde trafficking is compromised in fibroblasts from non-syndromic RP patients

Human dermal fibroblasts were obtained from a skin biopsy of three study subjects with disease associated *IFT140* variants: two with non-syndromic RP; homozygous c.998G>A (p.C333Y) and homozygous c.1451C>T (p.T484M) (Hull et al., 2016; Xu et al., 2015); and one individual with MSS and compound heterozygous variants c.932A>G (p.Y311C) and c.857-862delTTGA (p.I286Lfs*6) (Perrault et al., 2012) (**Figure 1A**). Analysis of fibroblasts by ultrastructure expansion microscopy (U-ExM) showed that IFT140 was enriched at the base of the cilium in controls and all patient lines, however the level of IFT140 was significantly reduced at the base of the cilium in all patient lines, particularly in the *IFT140^T484M/T484M^*line (**Figure 2A-C**). All patient lines showed enrichment of IFT88 at the tip of the cilium, indicative of deficient IFT retrograde transport (**Figure 2D-F**). Ciliation was not affected in any of the patient lines (**Figure 2G,H**). The cilia were significantly shorter in the *IFT140^T484M/T484M^* and *IFT140^Y311C/I286Lfs*6^* patient lines, but not in the *IFT140^C333Y/C333Y^* patient line (**Figure 2I**). The IFT-A interactor tubby-like protein 3 (TULP3) showed reduced fluorescence intensity within the cilium across all patient lines, but was not differentially enriched at either the base or tip, suggesting reduced but not fully disrupted trafficking (**Supplementary Figure 2**).

**Figure 2.**
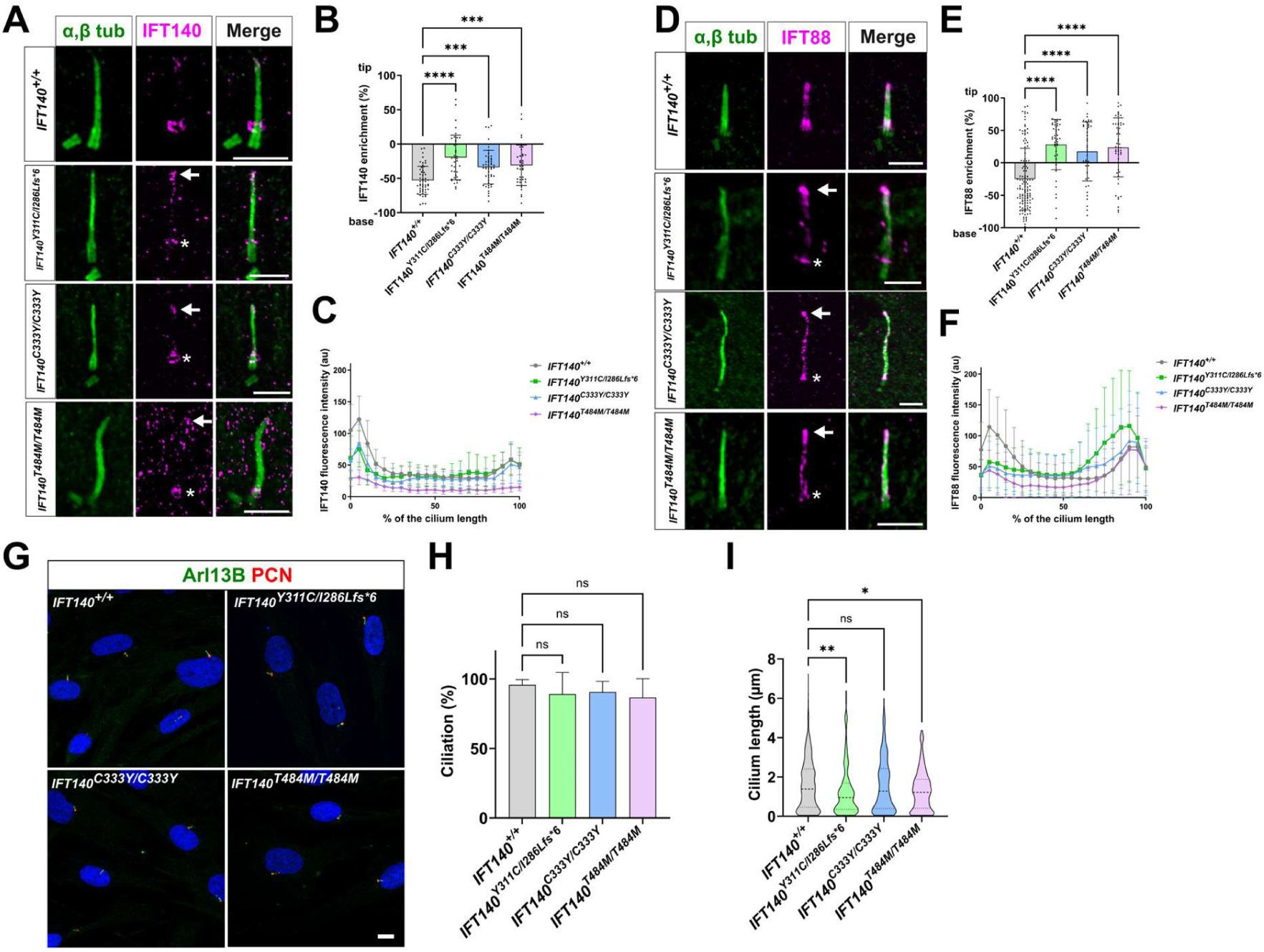
Expansion microscopy of patient fibroblasts reveals reduced levels of IFT140 at the base and accumulation of IFT88 at the cilia tip. **A)** Cilia were expanded up to 4.5 times and stained with IFT140 (magenta) and α, β tubulin (green). White arrows mark the tip of the cilium. Scale bar shows the size after expansion (5 µm). **B)** Percentage of the total IFT140 fluorescence enrichment at the ciliary tip is shown for each fibroblast cell line: *IFT140^+/+^*(n=51 cilia), *IFT140^Y311C/I286Lfs*6^* (n=40 cilia), *IFT140^C333Y/C333Y^* (n=51 cilia), *IFT140^T484M/T484M^* (n=39 cilia). Kruskal-Wallis test with Dunn’s multiple comparison test was used. ***p<0.001, ****p<0.0001. **C)** IFT140 fluorescence intensity measurement of IFT140 along the cilium length: *IFT140^+/+^* (n=51 cilia), *IFT140^Y311C/I286Lfs*6^* (n=40 cilia), *IFT140^C333Y/C333Y^* (n=51 cilia), *IFT140^T484M/T484M^* (n=39 cilia). **D)** Cilia were expanded up to 4.5 times and stained with IFT88 (magenta) and α, β tubulin (green). White arrows mark the tip of the cilium. Asterisk marks the base of the cilium. Scale bar shows the size after expansion (5 µm). **E)** Percentage of the total IFT88 fluorescence enrichment at the tip is shown for each fibroblast cell line. *IFT140^+/+^* (n=125 cilia), *IFT140^Y311C/I286Lfs*6^* (n=44 cilia), *IFT140^C333Y/C333Y^* (n=44 cilia), *IFT140^T484M/T484M^* (n=52 cilia). Kruskal-Wallis test with Dunn’s multiple comparison test. ****p<0.0001. **F)** Fluorescence intensity measurements of IFT88 along the cilium of fibroblasts: *IFT140^+/+^* (n=51 cilia), *IFT140^Y311C/I286Lfs*6^* (n=40 cilia), *IFT140^C333Y/C333Y^* (n=51 cilia), *IFT140^T484M/T484M^* (n=39 cilia). **G)** Representative image of the different fibroblast lines stained with the markers ARL13B (green) and PCN (red). Scale bar: 5 µm. **H)** Ciliation was measured for the presence of PCN with ARL13B positive staining. A minimum of four wells and at least 17 images were measured for each genotype. Kruskal-Wallis test with Dunn’s multiple comparison test was used. **I)** Cilium length was measured as the length of ARL13B using CiliaQ. The number of cilia counted are as follows: *IFT140^+/+^* (n=531 cilia), *IFT140^Y311C/I286Lfs*6^* (n=147 cilia), *IFT140^C333Y/C333Y^* (n=332 cilia), *IFT140^T484M/T484M^* (n=227 cilia). Kruskal-Wallis test with Dunn’s multiple comparison test. *p<0.05, **p<0.01.

### IFT140 RPE models have shorter cilia and accumulate IFT88 at the tip

To investigate the function of IFT140 in the retina, we generated a knock-out iPSC line of *IFT140* by CRISPR/Cas9 technology, resulting in the change c.T12del/c.T12ins (p.Y5Mfs*4/p.Y5Lfs*1) (*IFT140^KO^)*, (**Supplementary Figure 3**). In addition, we reprogrammed the *IFT140^T484M/T484M^* patient fibroblast line to obtain iPSCs (**Supplementary Figure 3**). The patient *IFT140^T484M/T484M^*, the *IFT140^KO^* and isogenic control lines were then differentiated to iPSC-RPE, using previously described protocols (Corral-Serrano et al., 2023; Michelet et al., 2020).

*IFT140^KO^* and *IFT140^T484M/T484M^* iPSC-RPE stained positive for PMEL17 showing presence of pigmentation in melanosomes, a key signature of RPE (**Figure 3A**). Expansion microscopy revealed IFT140 immunoreactivity was absent in the cilium of the *IFT140^KO^* line and barely detectable in the cilium of the *IFT140^T484M/T484M^* line (**Supplementary Figure 4**), consistent with the observations in the fibroblasts. There were no differences in IFT140 enrichment at either the base or tip, compared to the control.

**Figure 3.**
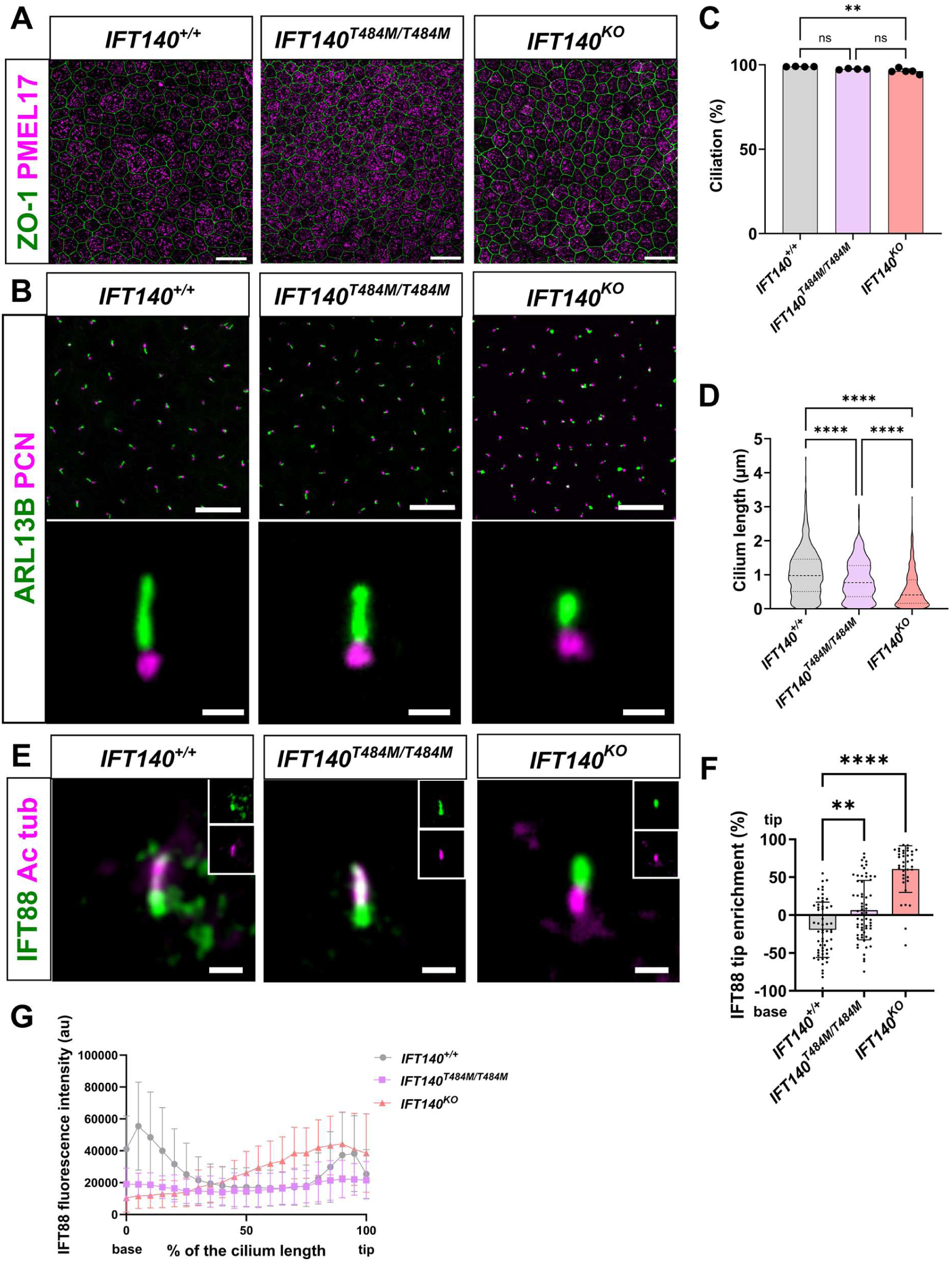
IFT140 iPSC-RPE show altered cilia formation and traffic defects. **A)** Immunostaining of the cell junction marker ZO-1 (green) and the melanosome marker PMEL17 (magenta) confirm RPE morphological appearance. Scale bar: 20 µm. **B)** iPSC-RPE were stained with ARL13B (green) and pericentrin (PCN, magenta). Scale bar: 20 µm upper panels, 1 µm lower panels. **C)** Ciliation was measured by the elaboration of ARL13B positive staining from a PCN positive basal body. A minimum of four wells from ≤3 independent differentiations were measured. Kruskal-Wallis test with Dunn’s multiple comparison test was used. ** P-value = 0.0096. **D)** Cilium length was measured using CiliaQ. A minimum of four wells from n≥3 independent differentiations were measured. The number of cilia counted are as follows: *IFT140^+/+^* (n=678 cilia), *IFT140^T484M/T484M^*(n=602 cilia), *IFT140^KO^* (n=1136 cilia). Kruskal-Wallis test with Dunn’s multiple comparison test was used. **** P-value < 0.0001. **E)** Immunostaining of IFT88 (green) and acetylated tubulin (Ac tub, magenta) in iPSC-RPE. Small insets on the top right show the channel separation. Scale bar: 1 µm. **F)** Enrichment of IFT88 at the cilia base or tip of *IFT140^+/+^* (n=57 cilia), *IFT140^T484M/T484M^*(n=62 cilia) and *IFT140^KO^* (n=37 cilia) iPSC-RPE. ** p< 0.01, ****, p<0.0001, based on Kruskal-Wallis test with Dunn’s multiple comparison test. **G)** Fluorescence intensity measurements of IFT88 along the cilium of iPSC-RPE in *IFT140^+/+^*(n=94), *IFT140^T484M/T484M^* (n=62) and *IFT140^KO^* (n=37).

The *IFT140^T484M/T484M^* line had shorter cilia than controls, while the *IFT140^KO^* line had shorter cilia than both the control and *IFT140^T484M/T484M^* line (**Figure 3B**). Ciliation incidence was not affected in the *IFT140^T484M/T484M^*line, and although there were significantly fewer cilia in the *IFT140^KO^*line, the ciliation percentage was still >95% (**Figure 3C**). The *IFT140^T484M/T484M^*iPSC-RPE showed differences in enrichment of IFT88 within the cilia axoneme and at the ciliary tip (**Figure 3E-G**), which was more pronounced in the *IFT140^KO^* iPSC-RPE, indicating deficient IFT retrograde transport (**Figure 3E-G**). Intriguingly, we observed that CEP290, a transition zone protein, was mislocalized in some of the cilia (10.63%, n=5/47 cilia) apical to the acetylated tubulin stain in the *IFT140^KO^* iPSC-RPE, suggesting altered ciliary gating (**Supplementary Figure 5**).

### Cilia structure and traffic are altered in IFT140 retinal organoid models

To investigate the function of IFT140 in human photoreceptors, retinal organoids (iPSC-ROs) were differentiated from *IFT140^KO^*iPSCs, their isogenic control, and the *IFT140^T484M/T484M^* patient line. At least 3 independent differentiations were carried out for each line over the course of several years. Immunofluorescence staining using ARL13B and PCN at day 90 of iPSC-ROs differentiation, before photoreceptor maturation, showed that the connecting cilium was shorter in both the *IFT140^T484M/T484M^* and *IFT140^KO^*line (**Supplementary Figure 6**). Immunofluorescence showed a reduction of IFT140 in the cilium of the *IFT140^T484M/T484M^* line and absence of IFT140 immunoreactivity in the *IFT140^KO^*line (**Supplementary Figure 7**).

The IFT-B complex protein, IFT88, and the bulge region protein LCA5, accumulated in the cilia of mature iPSC-ROs at day 200 of differentiation, showing increased staining distal to the basal body and extending into the axoneme, while the basal body marker pericentrin remained unchanged (**Figure 4A-G**). The alterations were most pronounced in the *IFT140^KO^* iPSC-ROs with the *IFT140^T484M/T484M^* iPSC-ROs intermediate between the knock-out and control iPSC-ROs. IFT88 had an average extension of 0.837 µm ± 0.034 µm (mean ± SEM) in the *IFT140^T484M/T484M^* iPSC-ROs and 1.759 µm ± 0.065 µm in *IFT140^KO^* iPSC-ROs. In the *IFT140^KO^* line, IFT88 even extended into the OS in some cilia (**Figure 4A**, inset), compared to the controls (0.691 µm ± 0.017 µm), (**Figure 4D**). LCA5 had a slight extension of 0.75 µm ± 0.030 µm in the *IFT140^T484M/T484M^* line compared to the controls (0.713 µm ± 0.030 µm), and 1.37 µm ± 0.058 in the *IFT140^KO^* line, suggesting disruption of the photoreceptor bulge region (**Figure 4E**). The transition zone protein CEP290 also had slightly altered localization, with an extension of 0.537 µm ± 0.011 in controls, 0.582 µm ± 0.012 µm in the *IFT140^T484M/T484M^* line, and 0.726 µm ± 0.015 µm in the *IFT140^KO^* line, indicating that the connecting cilium is mostly preserved (**Figure 4F**).

**Figure 4.**
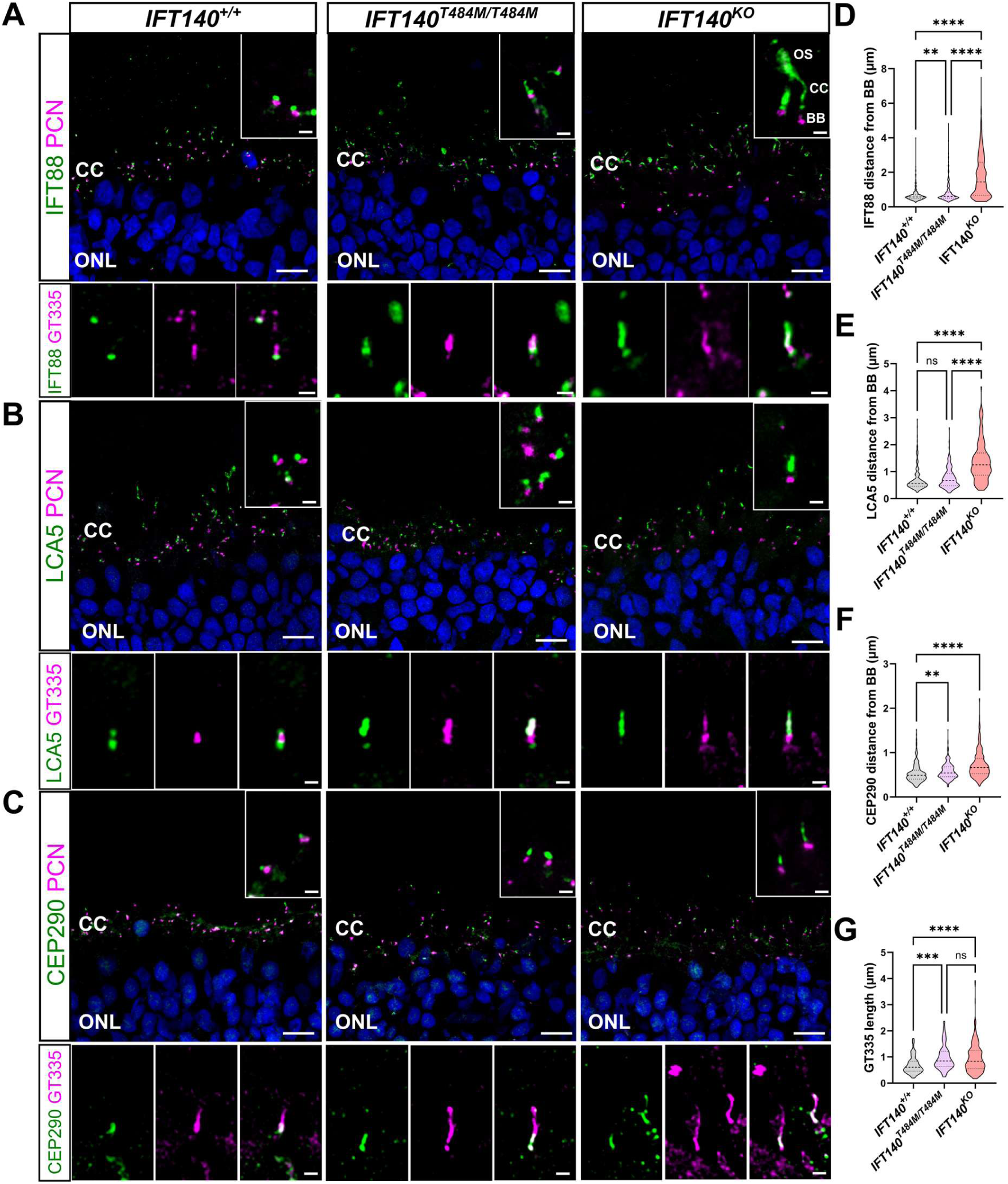
Cilia proteins accumulate along the photoreceptor connecting cilium and outer segments in IFT140 retinal organoids. iPSC-ROs from the *IFT140^KO^* line, isogenic controls, and the *IFT140^T484M/T484M^* patient line were collected at day 200 post differentiation. Data from ≥2 independent differentiations. **A)** Staining of the IFT complex B protein IFT88 (green) and PCN (magenta, top panel) or GT335 (magenta, bottom panel). Outer segment (OS), connecting cilia (CC), basal body (BB), outer nuclear layer (ONL). **B)** Staining for LCA5 (green) and PCN (magenta, top panel) or GT335 (magenta, bottom panel). **C)** Staining for CEP290 (green) and PCN (magenta, top panel) or GT335 (magenta, bottom panel). The inset shows a magnification of the connecting cilium (CC). Scale bar of low magnification images: 10 µm. Scale bar of insets: 1 µm. **D-G)** Quantification of **D)** the IFT88 distance from the basal body marked by the PCN staining in photoreceptor cilia of *IFT140^+/+^* (n=589 cilia, n=6 ROs), *IFT140^T484M/T484M^*, (n=359 cilia, n=4 ROs) and *IFT140^KO^* (n=358 cilia, n=5 ROs). **E)** Quantification of the LCA5 distance from the basal body marked by the PCN staining in photoreceptor cilia of *IFT140^+/+^* (n=205 cilia, n=6 ROs), *IFT140^T484M/T484M^* (n=154 cilia, n=4 ROs) and *IFT140^KO^*(n=153 cilia, n=5 ROs). **F)** Quantification of the CEP290 distance from the basal body marked by the PCN staining in photoreceptors cilia of control *IFT140^+/+^* (n=326 cilia, n=6 ROs), *IFT140^T484M/T484M^* (n=228 cilia, n=5 ROs) and *IFT140^KO^* (n=386 cilia, n=5 ROs). **G)** Quantification of the GT335 length in photoreceptor cilia of control *IFT140^+/+^* (n=106 cilia, n=6 ROs), *IFT140^T484M/T484M^* (n=54 cilia, n=4 ROs) and *IFT140^KO^* (n=228 cilia, n=5 ROs). All based on Kruskal-Wallis test with Dunn’s multiple comparison test, **p < 0.005, *** p-value < 0.001, **** p-value < 0.0001.

Co-staining with the polyglutamylation marker GT335 enabled visualization of the localization of these cilia proteins along the length of the connecting cilium (**Figure 4**). Interestingly, the pattern of GT335 staining extended further in the *IFT140^T484M/T484M^* patient line (0.963 ± 0.062 µm) and the *IFT140^KO^* line (0.964 µm ± 0.037 µm) compared to controls (0.685 µm ± 0.031) (**Figure 4G**), suggesting altered axonemal glutamylation. Collectively, these data support a model in which IFT140 dysfunction leads to defective photoreceptor retrograde trafficking, resulting in disruption of cilia compartmentalization and axonemal architecture in human photoreceptors.

Rhodopsin was used as functional readout of photoreceptor OS cilia traffic, as its delivery from the inner segment to the OS depends partially on intraflagellar transport (Awasthi et al., 2016; Bhowmick et al., 2009). Mislocalization of rhodopsin is a feature of photoreceptor ciliopathies and has been observed across multiple mouse models of retinal degeneration (Faber et al., 2023; Megaw et al., 2017; Rachel et al., 2015; Zhang et al., 2019) and iPSC-ROs (Athanasiou et al., 2025; Corral-Serrano et al., 2023; Sladen et al., 2024). Comparison of rhodopsin immunoreactivity fluorescence intensity in the OS to retention in the IS and outer nuclear layer (ONL) confirmed an accumulation of rhodopsin in the IS/ONL of photoreceptors in both *IFT140^T484M/T484M^* and *IFT140^KO^* lines (**Figure 5**). Analysis of L/M opsin staining did not reveal a significant accumulation in the IS/ONL in both *IFT140^T484M/T484M^* and *IFT140^KO^* lines compared to controls (**Supplementary Figure 8**).

**Figure 5.**
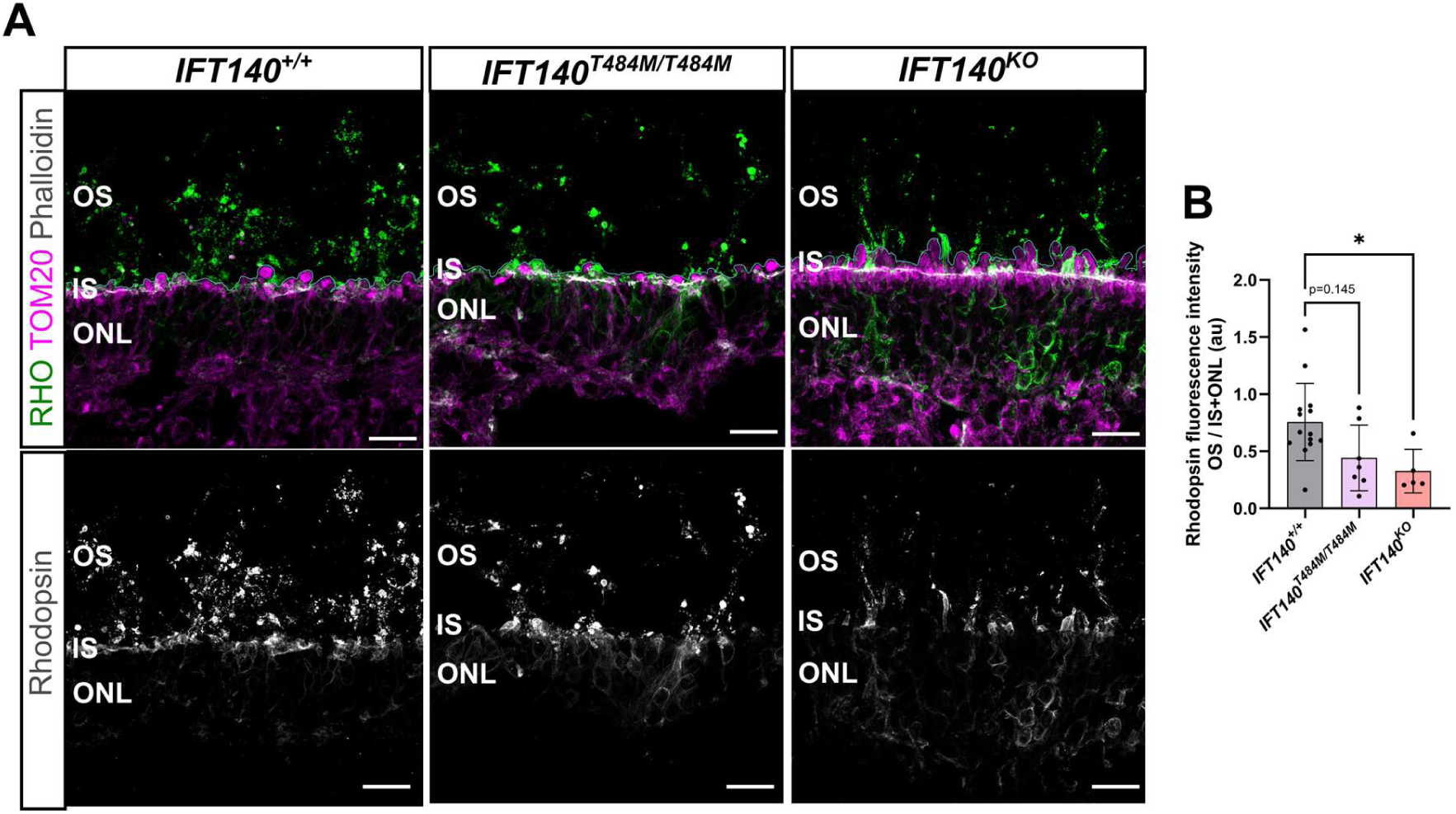
Rhodopsin accumulates in the IS and ONL in *IFT140^T484M/T484M^* and *IFT140^KO^* retinal organoids. iPSC-ROs from the *IFT140^KO^* line, isogenic controls *IFT140^+/+^*and the *IFT140^T484M/T484M^*patient line were collected at day 200 post differentiation, embedded and cryosectioned. Data from n=2 independent differentiations. **A)** Upper panel: staining of rhodopsin (green) and TOM20 (magenta) marking the inner segments. Phalloidin (grey) marks the outer limiting membrane. Lower panel: staining of rhodopsin (grey) only. Scale bar: 20 µm. **B)** The fluorescence intensity of rhodopsin was measured in the OS and the IS and ONL region (IS+ONL), and the ratio OS/IS+ONL derived. The analysis consisted of: *IFT140^+/+^*, n=14 images from n=7 ROs; *IFT140^T484M/T484M^*n=7 images from n=5 ROs; *IFT140^KO^*, n=5 images from n=3 ROs. Kruskal-Wallis test with Dunn’s multiple comparison test was used. p-value <0.05 (*).

### Cilia length and transport is improved in IFT140 RPE and retinal organoids following treatment with eupatilin

To evaluate the potential rescue of the observed phenotypes, a small molecule screen was performed in iPSC-RPE. Five different small molecules that have been reported to improve cilia defects in other ciliopathy models were used: the flavonoids eupatilin, quercetin and jaceosidin (Athanasiou et al., 2025; Corral-Serrano et al., 2023; Kim et al., 2018; Ortega et al., 2021; Wiegering et al., 2021); the rock2 inhibitor fasudil (Smith et al., 2025), and 2-IPMA (Choi et al., 2011). These compounds were tested on *IFT140^KO^*iPSC-RPE and compared to isogenic controls. Importantly, the flavonoids eupatilin, jaceosidin, and the combination of eupatilin with fasudil were able to improve cilia length in the *IFT140^KO^* line (**Supplementary Figure 9**). As a note, DMSO as vehicle had a small effect on cilium length in the *IFT140^KO^*line. Surprisingly, 2-IPMA had a negative effect on cilium length in the isogenic control and the *IFT140^KO^*line (**Supplementary Figure 9**). Fasudil or the combination of eupatilin 5 µM with fasudil 5 µM did not significantly improve cilium length in the *IFT140^KO^* or *IFT140^T484M/T484M^*lines (**Supplementary Figure 9**). Eupatilin treatment alone significantly improved cilium length in the *IFT140^T484M/T484M^* iPSC-RPE at 10 µM and 20 µM, while 10 µM eupatilin was the best concentration for the *IFT140^KO^* line (**Figure 6A, B**). Furthermore, 10 µM eupatilin was able to improve the IFT88 distribution within the cilium in both *IFT140^KO^* and *IFT140^T484M/T484M^* iPSC-RPE closer to that observed in controls (**Figure 6C-E**).

**Figure 6.**
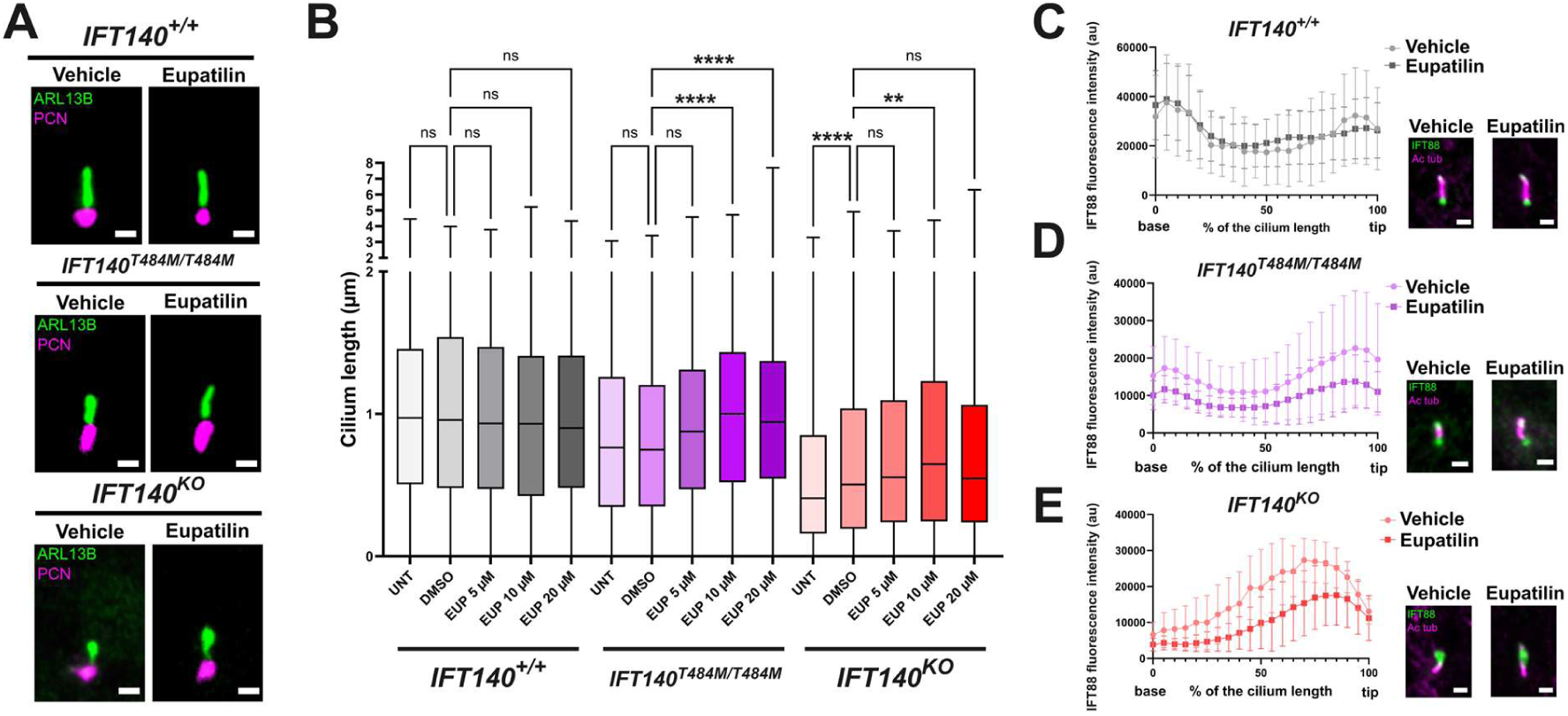
Eupatilin treatment improves cilia length and IFT88 traffic in IFT140 iPSC-RPE. **A)** Representative images of iPSC-RPE cilia from control, *IFT140^T484M/T484M^* and *IFT140^KO^*lines, stained with ARL13B (green) and PCN (magenta). Scale bar: 1 µm. **B)** Treatment of iPSCs-RPE with different concentrations of eupatilin (EUP) or vehicle (DMSO). UNT = untreated. Cells were treated for 24 hours with either EUP or DMSO and stained for ARL13B and PCN. Cilia length was measured using CiliaQ. At least 200 cilia were measured for each condition; ≥2 independent differentiations; and ≥3 independent treatments were performed for each condition. Significant differences between the treatment and the vehicle are shown as p<0.05 (*), p<0.01 (**), p<0.001 (***), p<0.0001 (****) based on Kruskal-Wallis test with Dunn’s multiple comparison test. **C-E)** IFT88 intensity fluorescence along the cilium in iPSC-RPE of ***B)*** *IFT140^+/+^*, **D**) *IFT140^T484M/T484M^* and **E**) *IFT140^KO^* treated with either DMSO or eupatilin 10 µM (EUP). The insets show a representative image of an iPSC-RPE cilium stained with IFT88 (green) and acetylated tubulin (ac tub, magenta).

To test whether eupatilin was able to rescue the defects observed in iPSC-ROs, all three lines were cultured until day 200 and then treated with 10 µM eupatilin or vehicle for an additional 30 days until day 230. The day 200 time point for treatment was selected as it corresponds to a stage at which organoids are more mature and exhibit more developed OS than at earlier time points. Eupatilin was able to reduce the IFT88 cilia accumulation in both lines, restoring the IFT88 distance from the basal body of the patient line to control levels (**Figure 7A** and **C**). Higher magnification of representative regions of the *IFT140^KO^* line suggests that, although IFT88 accumulation persists within the OS and is not fully cleared, its extension along the axoneme is markedly reduced (**Figure 7A** and **C**). In addition, rhodopsin trafficking to the OS and retention in the IS and ONL was improved post-treatment, although it did not reach significance for the *IFT140^T484M/T484M^* line (**Figure 7 B and D**). These data suggest restoration of retrograde transport in the disease retinal organoid models partially restores OS protein traffic following treatment with eupatilin.

**Figure 7.**
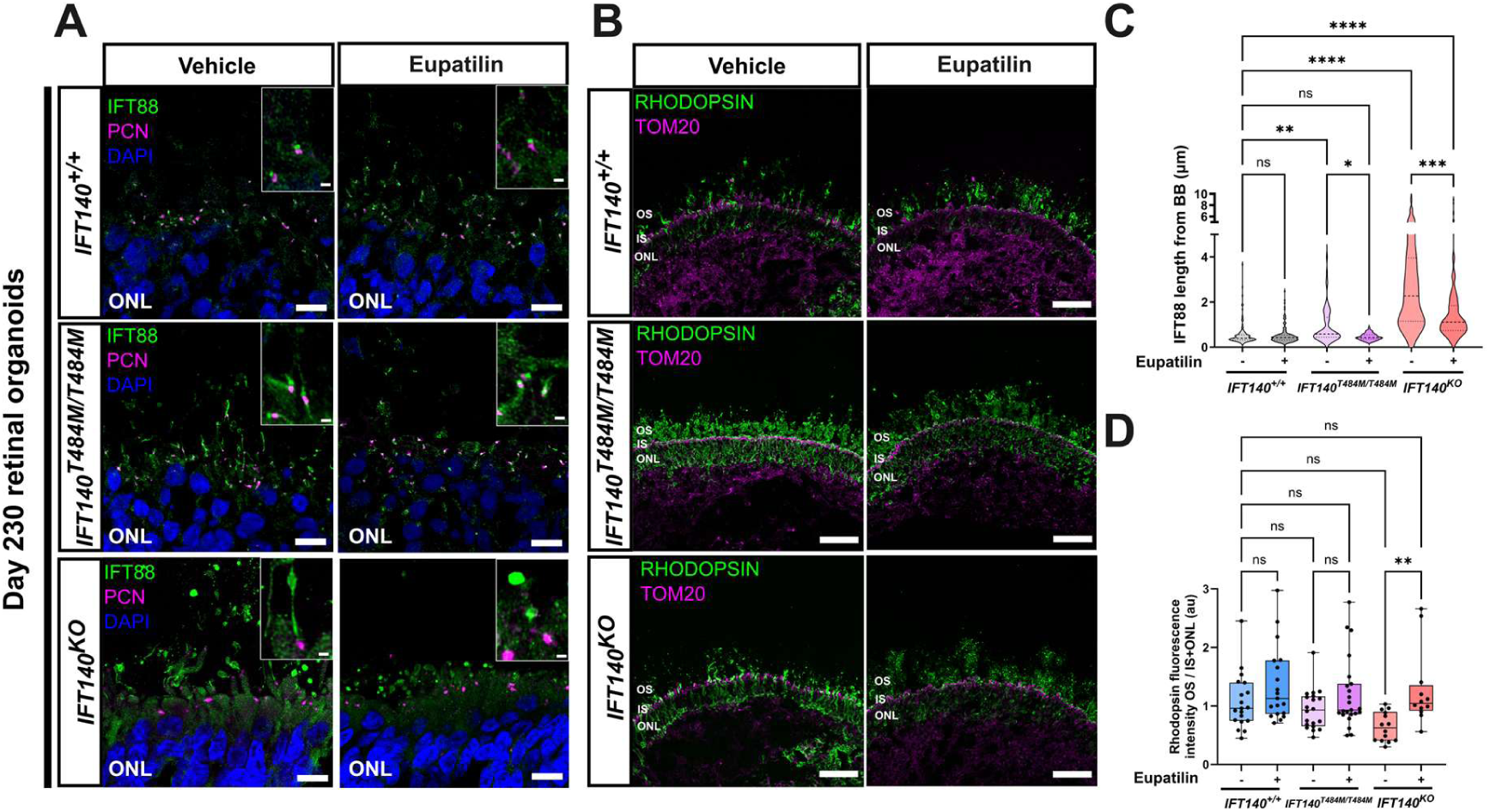
IFT88 and rhodopsin traffic is improved in IFT140 retinal organoids by eupatilin. iPSC-ROs from the *IFT140^KO^* line, isogenic controls, and the *IFT140^T484M/T484M^*patient line were differentiated for 200 days and then treated with 10 µM eupatilin or vehicle (DMSO) for further 30 days at day 200 post differentiation, embedded and cryosectioned. Data from n=2 independent differentiations. **A)** Immunostaining of the IFT- B protein IFT88 (green) and PCN (magenta). Scale bar of low magnification images: 10 µm. Scale bar of insets: 1 µm. **A)** Immunostaining of rhodopsin (green) and TOM20 (magenta) marking the IS for each line. Scale bar: 50 µm. **C)** Quantification of the IFT88 distance from the basal body marked by the PCN staining in photoreceptor cilia. The number of cilia measured were: *IFT140^+/+^*vehicle (n=123); *IFT140^+/+^* eupatilin (n=144); *IFT140^T484M/T484M^* vehicle (n=58); *IFT140^T484M/T484M^* eupatilin (n=53); *IFT140^KO^* vehicle (n=165); *IFT140^KO^* eupatilin (n=148). Kruskal-Wallis test with Dunn’s multiple comparison test of treated versus vehicle, * p < 0.05 (*), ** p < 0.01, *** p < 0.001, **** p < 0.0001. **D)** The fluorescence intensity of rhodopsin was measured in the OS and IS+ONL and the ratio OS/IS+ONL derived. The analysis consisted of: *IFT140^+/+^* vehicle (n=19 images from n=9 ROs); *IFT140^+/+^*eupatilin (n=19 images from n=7 ROs); *IFT140^T484M/T484M^* vehicle (n=20 images from n=5 ROs); *IFT140^T484M/T484M^* eupatilin (n=23 images from n=6 ROs); *IFT140^KO^* vehicle (n=14 images from n=6 ROs); *IFT140^KO^* eupatilin (n=12 images from n=6 ROs).

## Discussion

In this study, we generated and characterized novel models to investigate IFT140 associated retinal dystrophy, including mouse models, patient-derived fibroblasts and human iPSC-RPE and iPSC-ROs. The findings demonstrate that *IFT140* missense variants, or loss of IFT140, in human retinal cells results in impaired cilia traffic characterized by accumulation of IFT proteins at the cilia tip in RPE and along the photoreceptor connecting cilium and OS in iPSC-ROs, resulting in rhodopsin mislocalization. Importantly, knock-in mouse models carrying patient missense variants either had unexpected embryonic lethal phenotype, very different to the patient clinical presentation, or did not show overt retinal degeneration or dysfunction up to 12 months of age, suggesting potential species-specific differences in the impact of *IFT140* variants. This underscored the necessity for human stem cell-based systems to model the effects of retinal ciliopathy disease variants. In addition, we show that pharmacological treatment can improve the observed cilia phenotype in human retinal models.

A striking observation in this study is that the *Ift140* T484M variant has markedly different consequences in mice and humans. In mice, homozygous *Ift140^T484M/T484M^* animals were embryonic lethal, preventing assessment of retinal structure or function. This finding is consistent with the phenotype reported in the *cauli* mutant mice that has a homozygous missense mutation (c.2564T>A; p.I855K) in *Ift140*, which has not been identified as a pathogenic variant in humans to date (Miller et al., 2013). The homozygous *IFT140* T484M variant is associated with non-syndromic retinal dystrophy in humans without an overt syndromic phenotype (Hull et al., 2016). In contrast, homozygous *Ift140^Y311C/Y311C^* animals and compound heterozygous *Ift140^Y311C/T484M^* mice did not show any obvious structural changes indicative of retinal degeneration in the time frame examined. These findings indicate that the pathogenic consequences of *IFT140* missense variants are highly species- and context-dependent.

In the human fibroblasts, iPSC-RPE and iPSC-ROs we investigated, the T484M variant was associated with reduced IFT140 protein in the cilium together with accumulation of IFT88, consistent with impaired IFT-A function and defective retrograde trafficking. This suggests that the T484M variant is likely hypomorphic and could primarily affect IFT140 protein stability, rather than causing a complete loss of function in all tissues. We hypothesize that the threshold of IFT140 function required might differ between tissues with the retina particularly susceptible to lower levels caused by these variants.

At present, the basis for the difference in severity for the T484M variant between species remains unclear. One possibility is that this substitution in the mouse protein leads to a greater level of instability. This is supported by modelling of the structure of the human and mouse proteins that suggests the local environment could amplify the effects of the threonine to methionine change at residue 484 on protein instability. The human IFT140 amino acid residue 485 is proline, which would restrict any backbone movement resulting from the substitution, whereas in mouse this residue corresponds to serine, which could allow greater flexibility (**Supplementary Figure 10**). Modelling of the effect of the T484M substitution on the ΔΔG confirmed that it is predicted to destabilize IFT140 structure (>+1.00 kcal/mol), as does the Y311C substitution (+0.80 kcal/mol), but only T484M showed a difference between mouse and human with the mouse protein further destabilized by the T484M change (+0.25 kcal/mol) (**Supplementary Table S1**). This change in stability could lead to greater reductions in IFT140 levels in mouse than human. It is possible that the mouse embryo is more sensitive to reduced IFT140 function during early development, whereas in humans the higher protein levels associated with this variant permit embryonic survival but are insufficient to sustain long-term photoreceptor homeostasis.

Importantly, because homozygous *Ift140^T484M/T484M^* mice did not survive, we cannot exclude the possibility that they would also have developed a retinal phenotype had they lived long enough to develop a mature retina. Therefore, we cannot conclude that the T484M retina is unaffected in mice; rather, our data show that this allele causes a more severe developmental outcome in mice than in humans, while our data from our human stem cell-derived retinal models reveal a clear photoreceptor trafficking defect. These results point toward subtle dosage-sensitivity differences between species that determine disease outcomes. Together, these findings emphasize that mouse knock-in models may not faithfully predict the tissue-specific consequences of human *IFT140* missense variants, and highlight the value of human retinal ciliopathy models for mechanistic and therapeutic studies.

The connecting cilium represents a critical interface between the IS and OS compartments of photoreceptors, where coordinated IFT train turnaround is required to maintain compartmental integrity. Our data indicate that loss of IFT-A function disrupts this process, leading to impaired retrograde transport and defective recycling of ciliary components. This is evidenced by the pronounced distal accumulation and extension of IFT88, consistent with stalled anterograde trains that cannot be efficiently returned. The bulge-associated protein LCA5 also showed moderate distal extension, indicating disruption of the turnaround or bulge region, albeit to a lesser extent than IFT88. In contrast, the transition zone protein CEP290 exhibited only mild changes and did not extend into the outer segment, suggesting that proximal ciliary gating remains relatively preserved.

Similar alterations in compartmental organization at the connecting cilium tip and bulge region have been described in models of LCA5-associated retinopathy (Athanasiou et al., 2025; Faber et al., 2023), supporting the notion that disruption of IFT turnaround dynamics represents a shared pathogenic mechanism. Consistent with this, related phenotypes have been reported in *ift122* mutant zebrafish, where accumulation of IFT88 in the photoreceptor cilium and opsin mislocalization to the outer nuclear layer were observed (Boubakri et al., 2016), highlighting a conserved role for IFT-A components in maintaining photoreceptor compartmentalization.

The increased distal extension of GT335 staining in *IFT140^T484M/T484M^*and *IFT140^KO^* photoreceptors further indicates altered axonemal organization. This is reminiscent of recent work showing that imbalance in tubulin glutamylation disrupts photoreceptor ciliary architecture and leads to extension of CEP290 along the axoneme (Mercey et al., 2024). However, in contrast to the *Ccp5⁻/⁻* mouse model, where IFT88 levels are reduced, we observed mislocalization rather than depletion of IFT components. While total IFT88 protein levels were not directly quantified, the preservation of signal intensity together with its abnormal distal extension support the interpretation that the primary defect reflects mislocalization rather than depletion of IFT components, suggesting that altered glutamylation in IFT140 retinal dystrophy is unlikely to result from loss of CCP5 function. Instead, these changes may represent a secondary consequence of defective IFT train dynamics or impaired retrograde coordination.

Rhodopsin mislocalization was consistently observed in *IFT140^KO^*and *IFT140^T484M/T484M^* retinal organoids, whereas L/M opsin localization was comparatively preserved. This differential sensitivity suggests that rod photoreceptors may be particularly vulnerable to partial retrograde transport defects. These findings are consistent with the clinical presentation of *IFT140* mutations as RP, with earlier rod photoreceptor dysfunction, as opposed to compromised cone photoreceptor function.

Together, these findings support a model in which IFT140 deficiency disrupts IFT turnaround and boundary maintenance at the connecting cilium, leading to distal accumulation of ciliary proteins, altered axonemal organization, and defective outer segment protein trafficking. Further studies will be required to resolve the ultrastructural organization of the connecting cilium and the precise subciliary localization of these components consequent to IFT-A disruption.

A key finding of this study is that the flavonoid eupatilin improved cilium length and IFT88 distribution in both iPSC-derived RPE and iPSC-ROs, and partially rescued rhodopsin mislocalization and IFT88 accumulation in photoreceptors. These data extend previous reports of eupatilin-mediated rescue in other ciliopathy contexts (Athanasiou et al., 2025; Corral-Serrano et al., 2023; Kim et al., 2018; Tata et al., 2025; Wiegering et al., 2021) and demonstrate that pharmacological modulation of cilia traffic is feasible in human photoreceptors.

The precise mechanism of eupatilin action remains unclear. We have previously shown that eupatilin can reduce the expression of some cilia and phototransduction genes, including *RHO* (Corral-Serrano et al., 2023). The effect of eupatilin on IFT88 distribution suggests an improvement in IFT dynamics. This reduction may alleviate the trafficking burden and the load placed on the IFT machinery to correctly deliver, recycle, and remove proteins that is required at the connecting cilium, thereby facilitating more efficient transport under conditions of impaired IFT function. In support of this concept, downregulation of *Rho* expression has been recently shown to preserve photoreceptor integrity in mouse models of *Prph2* (Rutan Woods et al., 2024). The concurrent improvement in rhodopsin localization indicates that modulation of cilia transport can translate into correction of photoreceptor-specific cargo defects. Together, these findings support the concept of combining gene-agnostic therapeutic strategies targeting shared cilia transport pathways for the treatment of retinal ciliopathies.

## Materials and methods

### Animals

Missense mutations in mouse *Ift140* (GenBank accession number: NM_134126.3; Ensembl: ENSMUSG00000024169) were generated, as described (Beyer et al., 2024). Briefly, Cas9 mRNA (50 ng/μL), gRNA1 (12.5 ng/μL), gRNA2 (12.5 ng/μL), and donor oligo (100 ng/μL) were combined in 30 μL ddH20 and microinjected into one-cell stage zygotes. One-cell zygotes were transferred into pseudopregnant C57BL/6N females and progeny were genotyped by direct sequencing.

sgRNAs and donor oligos for homology-directed repair (HDR) were located in exons 8 and 12, for p.Y311C (TAC>TGC) and p.T484M (ACA>ATG), respectively. Forward (F) and reverse (R) sequences of sgRNAs that were used for each mutation are as follows: p.Y311C F: GAATTACATACTGAGTCTAC-<u>AGG</u>, R: CTGAGTCTACAGGAGAAGTT-<u>TGG</u>; p.T484M F: ATCAGTGCTAGCCATGCATG-<u>AGG</u>, R: CTAGCACTGATGTTTCACAC-<u>AGG</u>. (PAM sequences underlined).

For p.T484M 2 synonymous mutations p.V486V (GTG>GTA) and p.E491E (GAG>GAA) were also introduced into the donor oligo to prevent binding and re-cutting after HDR and the *ScaI* restriction site was used for genotyping (Primer-F: GAAGTAAGGCTGATAAGCAGAGT. Primer-R: TGAACTGACACGCATTTCCT. Wildtype allele: 612 bp. Mutant allele: 344 bp, 268 bp).

For p.Y311C 4 synonymous mutations p.I312I (ATA>ATC), p.L313L (CTG>TTA), p.L315L (CTA>CTC) and p.Q316Q (CAG>CAA) were also introduced into the donor oligo to prevent the binding and re-cutting of the sequence by gRNA after HDR and the *AflII* restriction site was used for genotyping (Primer-F: ATATGTAGCTGGGACTCCAAT. Primer-R: TTAAAACATAGACCAGGCTG. Wildtype allele: 501 bp. Mutant allele: 232 bp, 269 bp). Donor oligonucleotide sequences used can be found in **Supplementary Table S2**. These were crossed with C57BL/6J mice over several generations to remove the presence of the *rd8* mutation in *Crb1* in the C57BL/6N mice with screening for the rd8 allele (Mattapallil et al., 2012). Mouse genotyping primers can be found in **Supplementary Table S3**.

For mouse embryonic fibroblast isolation, mouse embryos at embryonic day 11.5 were minced, washed with PBS, and incubated with trypsin at 37°C for 30 minutes. Cells were collected by centrifugation and cultured in fibroblasts medium with primocin.

### ERG and OCT measurements

Mice were dark-adapted overnight and anaesthetized with Ketamine/Domitor (0.75 mL Ketamine, 0.5 mL Domitor, 0.75 mL sterile water at 0.1 mL/10 g). ERGs were carried out under red-light conditions. The scotopic activity of the retina was assessed as described before (Guarascio et al., 2025). A and B wave responses were analysed at log intensity 1 (log 10 cd*s/m2) by Excel (Microsoft) and Prism/GraphPad (Dotmatics).

Retinae were imaged using the Bioptigen Envisu R2300 Spectral-domain ophthalmic imaging system (SDOIS). To prevent cornea desiccation during the procedure, topical Systane Ultra (Alcon) lubricant eye drops were applied bilaterally. Following lubrication, mice were positioned for imaging in the animal imaging mount-rodent alignment stage (AIM-RAS). The position of the animals ensured that the images were centred to the optic nerve head. Images were acquired by the InVivoVue Clinic application using a rectangular scanning protocol consisting of a 1.4 mm by 1.4 mm perimeter with 750 A-scans (lines) and 10 B-scans (frames) with a 20 frames/B-scan. All OCT images were averaged and processed using the InVivoVue software. The Bioptigen InVivoVue Diver 2.0 was used to enable manual segmentation of the retinal layers and the outer nuclear layer thickness was measured after exporting results from Diver to Excel.

### Fibroblast cell lines

A skin biopsy was collected by Dr. Edward Bloch from a male patient carrying the variant c.1451C>T (p.T484M) in *IFT140* using a biopsy punch (Stiefel) and transferred to a 15 mL conical tube containing fibroblast medium (Dulbecco’s Modified Eagle’s Medium (DMEM)/F12, 10% Fetal Bovine Serum (FBS), 1% Penicillin–Streptomycin (Pen/Strep), 1% Sodium pyruvate). The skin biopsy was cultured in a gelatin-coated petri dish until fibroblasts started to appear. Control human dermal fibroblasts of male neonatal origin (HDFn) were obtained as previously described (Schwarz et al., 2017). To induce cilia growth, fibroblasts were incubated with serum starvation medium (DMEM/F12, 0.2% FBS, 1% Pen/Strep, 1% Sodium pyruvate).

### Ultrastructure expansion microscopy (U-ExM)

U-ExM of fibroblast cells was performed following a previously published protocol (Gambarotto et al., 2021). Cells were seeded onto 12mm round coverslips and grown to confluency, followed by overnight serum free starvation. Cells were then incubated overnight at 37 °C in anchoring solution consisting of 1.4% Formaldehyde (F8775, Sigma) and 2% Acrylamide (A4058, Sigma) prepared in 1x PBS. Following anchoring, a monomer solution was prepared by combining 25 μL of 40% Acrylamide (A4058, Sigma), 50 μL of 38% Sodium Acrylate (408220-25G, Sigma), 5 μL 2% N,N’-methylenbisacrylamide (M1533, Sigma) and 10 μL of 10x PBS (14200067, Gibco). This was supplemented with 5 μL of 10% Ammonium Persulfate (APS) and 5 μL 10% Tetra-methylethylenediamine (TEMED) to form the gelling solution. A 40 μL drop of gelling solution was placed onto parafilm and the coverslip with cell side facing down was placed on the droplet. Samples were incubated for 5 min at 4 °C, followed by 30 min at 37 °C to allow gelation. After polymerization, samples were gently removed from the parafilm. Each sample was incubated in 1 mL denaturation buffer (200 mM SDS, 200 mM NaCl, 50 mM Tris Base in water (pH 9)) in a 6 well plate for 10 min at RT with agitation, until the gel detached from the coverslip. Then the gel was subsequently transferred into a 1.5 mL Eppendorf tube filled with fresh denaturation buffer and incubated for 90 min at 95 °C. Following the denaturation step, gels were transferred to a 12 cm petri-dish containing ddH2O and incubated for 10 min. The water was replaced once, and the step was repeated to allow full expansion. The expansion factor was measured until the gel reached its maximum size. Gels were then washed with PBS for 5-10 min, cut in small square pieces, and transferred into 24 well plates.

For immunostaining, gels were incubated overnight at 4 °C with 500 μL of primary antibody diluted in 2% BSA/PBS. The following day, gels were washed three times with PBS containing 0.1% Tween-20 (PBST) for 10 min each at RT with agitation. Gels were incubated with 500 μl secondary antibody in 2% BSA/PBS for 1 hour at RT with agitation, followed by 3 additional washes in PBST. Prior to imaging, gels were transferred to a 6 well plate and incubated in ddH2O for at least 15 min to ensure full expansion. Gels were placed with cell side facing down onto 0.1mg/mL poly-D-Lysine-coated confocal dish (A3890401, Gibco). Gels were mounted in water with a round coverslip on the top of the gel to prevent drifting during imaging. Image acquisition was performed on an inverted confocal Leica Stellaris 8 microscope using 63x oil immersive objective with Lightning mode.

### Induced pluripotent stem cell (iPSCs) lines

*IFT140* c.1451C>T (p.T484M) patient fibroblasts were nucleofected using the Cell Line Nucleofector Kit R (Lonza) containing 1 μg of the episomal reprogramming vectors pCXLE-hOCT3/4-shp53-F (Addgene #27077), pCXLE-hUL Addgene #27080), pCXLE-hSK (Addgene #27078) and pSimple-miR302/367 (Addgene #98748), as previously described (Schwarz et al., 2015).

The control iPSCs used in this work were reprogrammed from healthy human dermal fibroblasts of male neonatal origin (HDFn) and were characterized previously (Corral-Serrano et al., 2025; Schwarz et al., 2015). The isogenic *IFT140^KO^*iPSC line to the HDFn line was generated using a simultaneous cellular reprogramming and CRISPR/Cas9 gene editing protocol (Howden et al., 2018), by targeting exon 1 in IFT140 in HDFn iPSCs. Control HDFn cells were nucleofected using the Cell Line Nucleofector Kit R (Lonza) containing 1 μg of each episomal reprogramming vector (Okita et al., 2011) and 1 μg of the IFT140-targeting PX458 plasmid. The antibodies used for characterization of iPSCs were the following: rabbit anti-OCT4 (Abcam) ab19857, 1:500; mouse anti-SSEA4 (Cell Signalling) 4755, 1:300; mouse anti-TRA-1-81 (Invitrogen) MA1-024, 1:1000.

### iPSCs differentiation to RPE

IPSCs were differentiated to RPE as previously described (Corral-Serrano et al., 2025). Briefly, iPSCs were seeded on Geltrex-coated (Thermo Fisher) six-well plates with mTESR Plus Medium (Stem Cell Technologies) until reaching full confluency. On day 0, cells were cultured with neural induction media (1:1 DMEM/F12/GlutaMAX™ (Gibco™), neurobasal medium (Gibco™), 55 μM 2-mercaptoethanol, 2% B27, 1% N2, 1% L-Glutamine and 1% Anti-Anti). On day 7, cells were cultured in RPE medium (DMEM, GlutaMAX™, 10% knock-out serum replacement (Gibco™), 1 x non-essential amino acids, 1 × L-Glutamine, 55 μM 2-mercaptoethanol (Gibco™), 1 x Anti-Anti. Fresh 50 ng/ml recombinant human activin A (PeproTech) was added with every media change. When the first signs of pigmentation became visible to the naked eye, activin A supplementation was stopped. Around day 40, iPSC-RPE were dissociated using Accutase (Sigma), filtered using a 40 μm cell strainer (Fisher Scientific), and transferred into a TC insert, for 24-well plates, PET, transparent, pore size 0.4 µm (Sarstedt).

For immunocytochemistry of iPSC-RPE, sections were first washed once in PBS, fixed in 2% PFA for 15 minutes, and blocked with 10% donkey serum, 0.05% Triton X-100 in PBS for 1 hour. Cells were then washed in PBS and incubated with primary antibodies. The primary antibodies used were the following: mouse anti-PMEL17 (Agilent, M0634) 1:50; rabbit anti-ZO1 (Thermo Fisher, 61-7300) 1:500. The primary antibodies were incubated in the blocking solution diluted 50% in PBS for 1 h. Sections were then washed in PBS and incubated with Alexa Fluor (Thermo Fisher) secondary antibodies in the diluted blocking solution for 45 min. Nuclei were stained using DAPI (2 μg/mL). Samples were washed in PBS and mounted using DAKO fluorescence mounting medium (Agilent).

For iPSC-RPE imaging of cilia, image acquisition was performed on an inverted confocal Leica Stellaris 8 microscope using 40x or 63x oil immersive objective with Lightning mode. For cilia length measurements, cilia were measured using CiliaQ (Hansen et al., 2021).

### iPSCs differentiation to iPSC-ROs

iPSCs were differentiated into iPSC-ROs as previously described (Corral-Serrano et al., 2020, 2023). Briefly, iPSCs were seeded on Geltrex-coated (Thermo Fisher) six-well plates with mTESR Plus Medium (Stem Cell Technologies) until 80-90% confluency. Essential 6™ Medium (Thermo Fisher) was added to the culture on Day 0 and Day 1, followed by the addition of neural induction media (Advanced DMEM/F-12 (1:1), 1% N2 supplement, 2 mM GlutaMax, and 1% Pen/Strep) on Day 2. Media was changed three times per week until neuro retinal vesicles (NRVs) appeared. On day 6 of differentiation, a single treatment with 1.5 nM BMP4 (Prepotech) was added, and half-media changes were carried out until day 16. NRVs were excised and kept in 25 square low attachment dishes for maturation. For retinal differentiation and maturation, serum-free retinal differentiation media [DMEM/F12 (3:1), 2% B27 (Thermo Fisher), 1% non-essential amino-acids (Thermo Fisher), and 1% Anti-Anti (Thermo Fisher)] was added for 6 days after collection. For retinal maturation, media was supplemented with 10% FBS (Gibco), 100 µM Taurine (Sigma-Aldrich), and 2 mM GlutaMax (Thermo Fisher). Retinoic acid (1 µM) was added on day 50. On day 70, N2 was added to the media (RMM2), and the concentration of retinoic acid was reduced to 0.5 µM. To promote photoreceptor differentiation, retinoic acid was removed from the media on day 100.

### Cryopreservation and immunohistochemistry of iPSC-ROs

IPSC-ROs were briefly washed once in PBS and placed in a bijou containing a mixture of 4% PFA + 5% sucrose in PBS for 40 min. IPSC-ROs were then placed in 6.25% sucrose in PBS for 1 h, followed by 12.5% sucrose in PBS for 1 h and in 25% sucrose in PBS for 1 h. All incubation steps were performed at 4°C. IPSC-ROs were then embedded in OCT, slowly frozen on dry ice, and stored at −80 °C until cryosectioning. For detection of cilia markers LCA5, GT335, IFT88, CEP290 and IFT140, iPSC-ROs were not fixed and briefly washed in PBS, before direct transfer to a block with OCT, slowly frozen on dry ice, and stored at −80 °C until cryosectioning. Cryosections (7-10 µm thick) were collected and stored at −20 °C for later analysis.

For immunohistochemistry of iPSC-ROs, sections were first washed once in PBS and blocked with 10% donkey serum, 0.05% Triton X-100 in PBS for 1 hour. For unfixed sections, sections were gently washed in PBS, fixed in 1% PFA for 15 minutes, washed with PBS, and permeabilized with 0.05% Tween20 in PBS for 15 minutes. The primary antibodies used were the following: rabbit anti-CEP290 (Abcam) ab84870, 1:100; rabbit anti-ARL13B (Proteintech) 17711-1-AP, 1:1000; mouse anti-Pericentrin (Abcam) ab28144, 1:1000; mouse anti-Rhodopsin 4D2 (Merck) MABN15, 1:1000; rabbit anti-Opsin Antibody, Red/Green (Merck) AB5405, 1:500; rabbit anti-IFT140 (Proteintech) 17460-1-AP to detect human IFT140, 1:200; rabbit anti-IFT140 (gift from Prof. Greg Pazour) to detect mouse IFT140, 1:500; rabbit anti-LCA5 (gift from Prof. Ronald Roepman), 1:100; mouse anti-GT335 (AG-20B-0020-C100), 1:1000; rabbit anti-IFT88 (Proteintech) 13967-1-AP, 1:200. The primary antibodies were incubated in the blocking solution diluted 50% in PBS for 1 h. Sections were then washed in PBS and incubated with Alexa Fluor (Thermo Fisher) secondary antibodies in the diluted blocking solution for 45 min. Nuclei were stained using DAPI (2 μg/mL). Samples were washed in PBS and mounted using DAKO fluorescence mounting medium (Agilent).

For iPSC-ROs imaging of cilia, image acquisition was performed on an inverted confocal Leica Stellaris 8 microscope using 63x oil immersive objective with Lightning mode. For iPSC-ROs imaging of rhodopsin staining, a 20x oil immersive objective was used. Images were processed using Fiji (Schindelin et al., 2012). Rhodopsin fluorescence intensity was measured on Fiji, using manually delineated lines to separate the OS from the IS+ONL region, as marked with a line in cyan (over the IS) in Figure 5.

## Acknowledgements

We thank Prof. Dr. Ronald Roepman for kindly providing the LCA5 antibody. We thank Prof. Dr. Greg Pazour for kindly providing the IFT140 antibody. We thank Dr. Katarina Jovanovic for her assistance in gene editing. We thank Mr. Edward Bloch for helping collect the skin biopsy. We also thank members of the Cheetham lab for their assistance with cell culture. We are grateful to the cilia research community for their valuable support and feedback. Finally, we are deeply grateful to the patients and their families for their participation in this study.

## Author contributions

Conceptualization: J.C.C.S., N.S., M.E.C.; Investigation: J.C.C.S., Y.J., N.S., S.G. R.G., K.Z., D.O., A.B. S.E.N., M.S., J.J.S., E.L.; Data analysis, J.C.C.S., Y.J., K.Z., A.B., R.G., S.E.N., Funding Acquisition, J.C.C.S., M.E.C., A.H., D.J.; Supervision: J.C.C.S., M.E.C., D.J.; Visualization: J.C.C.S, Y.J.; Writing - original draft: J.C.C.S., M.E.C.; Writing - review and editing, all authors.

## References

Absalon, S., Blisnick, T., Kohl, L., Toutirais, G., Doré, G., Julkowska, D., Tavenet, A., & Bastin, P. (2008). Intraflagellar Transport and Functional Analysis of Genes Required for Flagellum Formation in Trypanosomes. Molecular Biology of the Cell, 19(3), 929–944. 10.1091/mbc.E07-08-0749

Athanasiou, D., Afanasyeva, T. A. V., Chai, N., Ziaka, K., Jovanovic, K., Guarascio, R., Boldt, K., Corral-Serrano, J. C., Kanuga, N., Roepman, R., Collin, R. W. J., & Cheetham, M. E. (2025). Small molecule treatment alleviates photoreceptor cilia defects in LCA5-deficient human retinal organoids. Acta Neuropathologica Communications, 13(1), 26. 10.1186/s40478-025-01943-y

Awasthi, M., Ranjan, P., Sharma, K., Veetil, S. K., & Kateriya, S. (2016). The trafficking of bacterial type rhodopsins into the Chlamydomonas eyespot and flagella is IFT mediated. Scientific Reports, 6(1), 34646. 10.1038/srep34646

Beyer, T., Diwan, G. D., Leonhard, T., Dahlke, K., Klose, F., Stehle, I. F., Seda, M., Bolz, S., Woerz, F., Russell, R. B., Jenkins, D., Ueffing, M., & Boldt, K. (2025). Ciliopathy-Associated Missense Mutations in IFT140 are Tolerated by the Inherent Resilience of the IFT Machinery. Molecular & Cellular Proteomics: MCP, 24(3), 100916. 10.1016/j.mcpro.2025.100916

Beyer, T., Martins, T., Srikaran, J. J., Seda, M., Peskett, E., Klose, F., Junger, K., Beales, P. L., Ueffing, M., Boldt, K., & Jenkins, D. (2024). Affinity Purification of Intraflagellar Transport (IFT) Proteins in Mice Using Endogenous Streptavidin/FLAG Tags. In V. Mennella (Ed.), Cilia: Methods and Protocols (pp. 199–212). Springer US. 10.1007/978-1-0716-3507-0_12

Bhowmick, R., Li, M., Sun, J., Baker, S. A., Insinna, C., & Besharse, J. C. (2009). Photoreceptor IFT complexes containing chaperones, guanylyl cyclase 1 and rhodopsin. Traffic, 10(6), 648–663. 10.1111/j.1600-0854.2009.00896.x

Boubakri, M., Chaya, T., Hirata, H., Kajimura, N., Kuwahara, R., Ueno, A., Malicki, J., Furukawa, T., & Omori, Y. (2016). Loss of ift122, a Retrograde Intraflagellar Transport (IFT) Complex Component, Leads to Slow, Progressive Photoreceptor Degeneration Due to Inefficient Opsin Transport*. Journal of Biological Chemistry, 291(47), 24465–24474. 10.1074/jbc.M116.738658

Bujakowska, K. M., Liu, Q., & Pierce, E. A. (2017). Photoreceptor Cilia and Retinal Ciliopathies. Cold Spring Harbor Perspectives in Biology, 9(10). 10.1101/CSHPERSPECT.A028274

Corral-Serrano, J. C., Lamers, I. J. C., Van Reeuwijk, J., Duijkers, L., Hoogendoorn, A. D. M., Yildirim, A., Argyrou, N., Ruigrok, R. A. A., Letteboer, S. J. F., Butcher, R., Van Essen, M. D., Sakami, S., Van Beersum, S. E. C., Palczewski, K., Cheetham, M. E., Liu, Q., Boldt, K., Wolfrum, U., Ueffing, M., … Collin, R. W. J. (2020). PCARE and WASF3 regulate ciliary F-actin assembly that is required for the initiation of photoreceptor outer segment disk formation. Proceedings of the National Academy of Sciences of the United States of America, 117(18), 9922–9931. 10.1073/pnas.1903125117

Corral-Serrano, J. C., Sladen, P. E., Ottaviani, D., Rezek, O. F., Athanasiou, D., Jovanovic, K., van der Spuy, J., Mansfield, B. C., & Cheetham, M. E. (2023). Eupatilin Improves Cilia Defects in Human CEP290 Ciliopathy Models. Cells, 12(12), 1575. 10.3390/CELLS12121575/S1

Corral-Serrano, J. C., Vaclavik, V., Van de Sompele, S., Kaminska, K., Jovanovic, K., Escher, P., Van den Broeck, F., Cancellieri, F., Toulis, V., Leroy, B. P., de Zaeytijd, J., You, Z., Ottaviani, D., Quinodoz, M., Bordeanu, G., Hardcastle, A. J., Coppieters, F., Tran, V. H., Cheetham, M. E., … De Baere, E. (2025). A novel recurrent ARL3 variant c.209G > A p.(Gly70Glu) causes variable non-syndromic dominant retinal dystrophy with defective lipidated protein transport in human retinal stem cell models. Human Molecular Genetics, 34(9), 821–834. 10.1093/hmg/ddaf029

Dordoni, C., Zeni, L., Toso, D., Mazza, C., Mescia, F., Cortinovis, R., Econimo, L., Savoldi, G., Alberici, F., Scolari, F., & Izzi, C. (2024). Monoallelic pathogenic IFT140 variants are a common cause of autosomal dominant polycystic kidney disease-spectrum phenotype. Clinical Kidney Journal, 17(2), sfae026. 10.1093/ckj/sfae026

Faber, S., Mercey, O., Junger, K., Garanto, A., May-Simera, H., Ueffing, M., Collin, R. W. J., Boldt, K., Guichard, P., Hamel, V., & Roepman, R. (2023). Gene augmentation of *LCA5*-associated Leber congenital amaurosis ameliorates bulge region defects of the photoreceptor ciliary axoneme. JCI Insight, 8(10). 10.1172/jci.insight.169162

Francis, R., San Agustin, J. T., Szabo Rogers, H. L., Cui, C., Jonassen, J. A., Eguether, T., Follit, J. A., Lo, C. W., & Pazour, G. J. (2023). Autonomous and non-cell autonomous etiology of ciliopathy associated structural birth defects. bioRxiv : The Preprint Server for Biology. 10.1101/2023.06.07.544132

Fujimaru, T., Mori, T., Sekine, A., Chiga, M., Mandai, S., Kikuchi, H., Mori, Y., Hara, Y., Fujiki, T., Ando, F., Susa, K., Iimori, S., Naito, S., Hanazawa, R., Hirakawa, A., Mochizuki, T., Suwabe, T., Ubara, Y., Uchida, S., & Sohara, E. (2024). Importance of *IFT140* in Patients with Polycystic Kidney Disease Without a Family History. Kidney International Reports, 9(9), 2685–2694. 10.1016/j.ekir.2024.06.021

Gambarotto, D., Hamel, V., & Guichard, P. (2021). Ultrastructure expansion microscopy (U-ExM). Methods in Cell Biology, 161, 57–81. 10.1016/BS.MCB.2020.05.006

Griffiths, J. D., Ehidiamhen, G., Lopez-Garcia, S. C., Hubbard, R., Cook, J., & Ong, A. C. M. (2026). Monoallelic *IFT140* Variants Causing Childhood-Onset Autosomal Dominant Polycystic Kidney Disease. American Journal of Kidney Diseases, 87(1), 124–128. 10.1053/j.ajkd.2025.08.006

Guarascio, R., Ziaka, K., Hau, K.-L., Piccolo, D., Nieuwenhuis, S. E., Bakoulina, A., Asfahani, R., Aguilà, M., Athanasiou, D., Sefic Svara, D., Li, Y., Chen, R., & Cheetham, M. E. (2025). Preventing light-induced toxicity in a new mouse model of sector retinitis pigmentosa caused by Rhodopsin M39R variant. Cell Death Discovery, 11(1), 477. 10.1038/s41420-025-02769-2

Gupta, M., & Pazour, G. J. (2024). Intraflagellar Transport: A Critical Player in Photoreceptor Development and the Pathogenesis of Retinal Degenerative Diseases. *Cytoskeleton (Hoboken*, N.J.), 81(11), 556–568. 10.1002/cm.21823

Hansen, J. N., Rassmann, S., Stüven, B., Jurisch-Yaksi, N., & Wachten, D. (2021). CiliaQ: a simple, open-source software for automated quantification of ciliary morphology and fluorescence in 2D, 3D, and 4D images. The European Physical Journal. E, Soft Matter, 44(2). 10.1140/EPJE/S10189-021-00031-Y

Hesketh, S. J., Mukhopadhyay, A. G., Nakamura, D., Toropova, K., & Roberts, A. J. (2022). IFT-A structure reveals carriages for membrane protein transport into cilia. Cell, 185(26), 4971–4985.e16. 10.1016/J.CELL.2022.11.010

Hirano, T., Katoh, Y., Nakayama, K., & Marshall, W. (2017). Intraflagellar transport—A complex mediates ciliary entry and retrograde trafficking of ciliary G protein-coupled receptors. Molecular Biology of the Cell, 28(3), 429–439. 10.1091/mbc.E16-11-0813

Howden, S. E., Thomson, J. A., & Little, M. H. (2018). Simultaneous reprogramming and gene editing of human fibroblasts. Nature Protocols 2018 13:5, 13(5), 875–898. 10.1038/nprot.2018.007

Hull, S., Owen, N., Islam, F., Tracey-White, D., Plagnol, V., Holder, G. E., Michaelides, M., Carss, K., Raymond, F. L., Rozet, J. M., Ramsden, S. C., Black, G. C. M., Perrault, I., Sarkar, A., Moosajee, M., Webster, A. R., Arno, G., & Moore, A. T. (2016). Nonsyndromic retinal dystrophy due to bi-allelic mutations in the ciliary transport gene IFT140. Investigative Ophthalmology and Visual Science, 57(3), 1053–1062. 10.1167/iovs.15-17976

Jiang, M., Palicharla, V. R., Miller, D., Hwang, S.-H., Zhu, H., Hixson, P., Mukhopadhyay, S., & Sun, J. (2023). Human IFT-A complex structures provide molecular insights into ciliary transport. Cell Research, 33(4), 288–298. 10.1038/s41422-023-00778-3

Jonassen, J. A., SanAgustin, J., Baker, S. P., & Pazour, G. J. (2012). Disruption of IFT Complex A Causes Cystic Kidneys without Mitotic Spindle Misorientation. Journal of the American Society of Nephrology, 23(4), 641. 10.1681/ASN.2011080829

Kim, Y. J., Kim, S., Jung, Y., Jung, E., Kwon, H. J., & Kim, J. (2018). Eupatilin rescues ciliary transition zone defects to ameliorate ciliopathy-related phenotypes. Journal of Clinical Investigation, 128(8), 3642–3648. 10.1172/JCI99232

Lacey, S. E., Foster, H. E., & Pigino, G. (2023). The molecular structure of IFT-A and IFT-B in anterograde intraflagellar transport trains. Nature Structural & Molecular Biology, 30(5), 584–593. 10.1038/s41594-022-00905-5

Lacey, S. E., Graziadei, A., & Pigino, G. (2024). Extensive structural rearrangement of intraflagellar transport trains underpins bidirectional cargo transport. Cell, 187(17), 4621–4636.e18. 10.1016/j.cell.2024.06.041

Mattapallil, M. J., Wawrousek, E. F., Chan, C.-C., Zhao, H., Roychoudhury, J., Ferguson, T. A., & Caspi, R. R. (2012). The Rd8 Mutation of the Crb1 Gene Is Present in Vendor Lines of C57BL/6N Mice and Embryonic Stem Cells, and Confounds Ocular Induced Mutant Phenotypes. Investigative Ophthalmology & Visual Science, 53(6), 2921–2927. 10.1167/iovs.12-9662

Megaw, R., Abu-Arafeh, H., Jungnickel, M., Mellough, C., Gurniak, C., Witke, W., Zhang, W., Khanna, H., Mill, P., Dhillon, B., Wright, A. F., Lako, M., & Ffrench-Constant, C. (2017). Gelsolin dysfunction causes photoreceptor loss in induced pluripotent cell and animal retinitis pigmentosa models. Nature Communications 2017 8:1, 8(1), 1–10. 10.1038/s41467-017-00111-8

Meleppattu, S., Zhou, H., Dai, J., Gui, M., & Brown, A. (2022). Mechanism of IFT-A polymerization into trains for ciliary transport. Cell, 185(26), 4986–4998.e12. 10.1016/j.cell.2022.11.033

Mercey, O., Gadadhar, S., Magiera, M. M., Lebrun, L., Kostic, C., Moulin, A., Arsenijevic, Y., Janke, C., Guichard, P., & Hamel, V. (2024). Glutamylation imbalance impairs the molecular architecture of the photoreceptor cilium. The EMBO Journal, 43(24), 6679–6704. 10.1038/s44318-024-00284-1

Mill, P., Christensen, S. T., & Pedersen, L. B. (2023). Primary cilia as dynamic and diverse signalling hubs in development and disease. Nature Reviews Genetics, 24(7), 421– 441. 10.1038/s41576-023-00587-9

Miller, K. A., Ah-Cann, C. J., Welfare, M. F., Tan, T. Y., Pope, K., Caruana, G., Freckmann, M.-L., Savarirayan, R., Bertram, J. F., Dobbie, M. S., Bateman, J. F., & Farlie, P. G. (2013). Cauli: A Mouse Strain with an Ift140 Mutation That Results in a Skeletal Ciliopathy Modelling Jeune Syndrome. PLoS Genetics, 9(8), e1003746. 10.1371/journal.pgen.1003746

Nachury, M. V. (2022). The gymnastics of intraflagellar transport complexes keeps trains running inside cilia. Cell, 185(26), 4863–4865. 10.1016/j.cell.2022.12.005

Okita, K., Matsumura, Y., Sato, Y., Okada, A., Morizane, A., Okamoto, S., Hong, H., Nakagawa, M., Tanabe, K., Tezuka, K. I., Shibata, T., Kunisada, T., Takahashi, M., Takahashi, J., Saji, H., & Yamanaka, S. (2011). A more efficient method to generate integration-free human iPS cells. Nature Methods 2011 8:5, 8(5), 409–412. 10.1038/nmeth.1591

Ortega, J. T., Parmar, T., Golczak, M., & Jastrzebska, B. (2021). Protective effects of flavonoids in acute models of light-induced retinal degeneration. Molecular Pharmacology, 99(1), 60–77. 10.1124/MOLPHARM.120.000072/-/DC1

Perrault, I., Saunier, S., Hanein, S., Filhol, E., Bizet, A. A., Collins, F., Salih, M. A. M., Gerber, S., Delphin, N., Bigot, K., Orssaud, C., Silva, E., Baudouin, V., Oud, M. M., Shannon, N., Le Merrer, M., Roche, O., Pietrement, C., Goumid, J., … Rozet, J. M. (2012). Mainzer-saldino syndrome is a ciliopathy caused by IFT140 mutations. American Journal of Human Genetics, 90(5), 864–870. 10.1016/j.ajhg.2012.03.006

Rachel, R. A., Yamamoto, E. A., Dewanjee, M. K., May-Simera, H. L., Sergeev, Y. V., Hackett, A. N., Pohida, K., Munasinghe, J., Gotoh, N., Wickstead, B., Fariss, R. N., Dong, L., Li, T., & Swaroop, A. (2015). CEP290 alleles in mice disrupt tissue-specific cilia biogenesis and recapitulate features of syndromic ciliopathies. Human Molecular Genetics, 24(13), 3775–3791. 10.1093/hmg/ddv123

Rivolta, C., Celik, E., Kamdar, D., Cancellieri, F., Kaminska, K., Ullah, M., Barberán-Martínez, P., Bouckaert, M., Cortón, M., Delanote, E., Fernández-Caballero, L., García, G. G., Holtes, L. K., Karali, M., Lopez, I., Peter, V. G., Schneider, N., Vincke, L., Ayuso, C., … Quinodoz, M. (2025). RetiGene, a comprehensive gene atlas for inherited retinal diseases. The American Journal of Human Genetics, 112(10), 2253–2265. 10.1016/j.ajhg.2025.08.017

Rutan Woods, C. T., Makia, M. S., Lewis, T. R., Crane, R., Zeibak, S., Yu, P., Kakakhel, M., Castillo, C. M., Arshavsky, V. Y., Naash, M. I., & Al-Ubaidi, M. R. (2024). Downregulation of rhodopsin is an effective therapeutic strategy in ameliorating peripherin-2-associated inherited retinal disorders. Nature Communications, 15(1), 4756. 10.1038/s41467-024-48846-5

Salhi, S., Doreille, A., Dancer, M. S., Boueilh, A., Filipozzi, P., El Karoui, K., Ponce, F., Lebre, A.-S., Raymond, L., & Mesnard, L. (2024). Monoallelic Loss-of-Function *IFT140* Pathogenic Variants Cause Autosomal Dominant Polycystic Kidney Disease: A Confirmatory Study With Suspicion of an Additional Cardiac Phenotype. American Journal of Kidney Diseases, 83(5), 688–691. 10.1053/j.ajkd.2023.08.019

Schindelin, J., Arganda-Carreras, I., Frise, E., Kaynig, V., Longair, M., Pietzsch, T., Preibisch, S., Rueden, C., Saalfeld, S., Schmid, B., Tinevez, J.-Y., White, D. J., Hartenstein, V., Eliceiri, K., Tomancak, P., & Cardona, A. (2012). Fiji: An open-source platform for biological-image analysis. Nature Methods, 9(7), 676–682. 10.1038/nmeth.2019

Schwarz, N., Carr, A. J., Lane, A., Moeller, F., Chen, L. L., Aguilà, M., Nommiste, B., Muthiah, M. N., Kanuga, N., Wolfrum, U., Nagel-wolfrum, K., Da cruz, L., Coffey, P. J., Cheetham, M. E., & Hardcastle, A. J. (2015). Translational read-through of the RP2 Arg120stop mutation in patient iPSC-derived retinal pigment epithelium cells. Human Molecular Genetics, 24(4), 972–986. 10.1093/hmg/ddu509

Schwarz, N., Lane, A., Jovanovic, K., Parfitt, D. A., Aguila, M., Thompson, C. L., da Cruz, L., Coffey, P. J., Chapple, J. P., Hardcastle, A. J., & Cheetham, M. E. (2017). Arl3 and RP2 regulate the trafficking of ciliary tip kinesins. Human Molecular Genetics, 26(17), 3451. 10.1093/hmg/ddx245

Seeman, T., Šuláková, T., Bosáková, A., Indráková, J., & Grečmalová, D. (2024). The First Pediatric Case of an IFT140 Heterozygous Deletion Causing Autosomal Dominant Polycystic Kidney Disease: Case Report. Case Reports in Nephrology and Dialysis, 14(1), 104–109. 10.1159/000539176

Senum, S. R., Li, Y. (Sabrina) M., Benson, K. A., Joli, G., Olinger, E., Lavu, S., Madsen, C. D., Gregory, A. V., Neatu, R., Kline, T. L., Audrézet, M. P., Outeda, P., Nau, C. B., Meijer, E., Ali, H., Steinman, T. I., Mrug, M., Phelan, P. J., Watnick, T. J., … Harris, P. C. (2022). Monoallelic IFT140 pathogenic variants are an important cause of the autosomal dominant polycystic kidney-spectrum phenotype. The American Journal of Human Genetics, 109(1), 136–156. 10.1016/J.AJHG.2021.11.016

Sladen, P. E., Naeem, A., Adefila-Ideozu, T., Vermeule, T., Busson, S. L., Michaelides, M., Naylor, S., Forbes, A., Lane, A., & Georgiadis, A. (2024). AAV-RPGR Gene Therapy Rescues Opsin Mislocalisation in a Human Retinal Organoid Model of RPGR- Associated X-Linked Retinitis Pigmentosa. International Journal of Molecular Sciences, 25(3). 10.3390/ijms25031839

Smith, C. E. L., Streets, A. J., Lake, A. V. R., Natarajan, S., Best, S. K., Szymanska, K., Karwatka, M., Stevenson, T., Trowbridge, R., Grant, G., Grellscheid, S. N., Foster, R., Morrison, C. G., Mavria, G., Bond, J., Ong, A. C. M., & Johnson, C. A. (2025). Drug and siRNA screens identify ROCK2 as a therapeutic target for ciliopathies. Communications Medicine, 5(1), 129. 10.1038/s43856-025-00847-1

Tata, A., Rocha, G., Hureaux, M., Serafin, A. S., Porée, E., Menguy, L., Goudin, N., Cagnard, N., Gréau, L., Fila, M., Briseño-Roa, L., Annereau, J.-P., Saunier, S., & Benmerah, A. (2025). Prostaglandin Analogs and Eupatilin as Treatments for Nephronophthisis. Kidney International Reports, 10(8), 2821–2835. 10.1016/j.ekir.2025.04.060

Wiegering, A., Dildrop, R., Vesque, C., Khanna, H., Schneider-Maunoury, S., & Gerhardt, C. (2021). Rpgrip1l controls ciliary gating by ensuring the proper amount of Cep290 at the vertebrate transition zone. Molecular Biology of the Cell, 32(8), 675–689. 10.1091/MBC.E20-03-0190

Xu, M., Yang, L., Wang, F., Li, H., Wang, X., Wang, W., Ge, Z., Wang, K., Zhao, L., Li, H., Li, Y., Sui, R., & Chen, R. (2015). Mutations in human IFT140 cause non-syndromic retinal degeneration. Human Genetics, 134(10), 1069. 10.1007/S00439-015-1586-X

Zhang, X., Shahani, U., Reilly, J., & Shu, X. (2019). Disease mechanisms and neuroprotection by tauroursodeoxycholic acid in Rpgr knockout mice. Journal of Cellular Physiology, 234(10), 18801–18812. 10.1002/jcp.28519

