## Supplementary Data for "Pharmacological rescue of cilia trafficking defects in IFT140 retinal organoid and RPE models of retinal dystrophy"

**Supplementary Figures and Supplementary Tables**

### Supplementary Figures

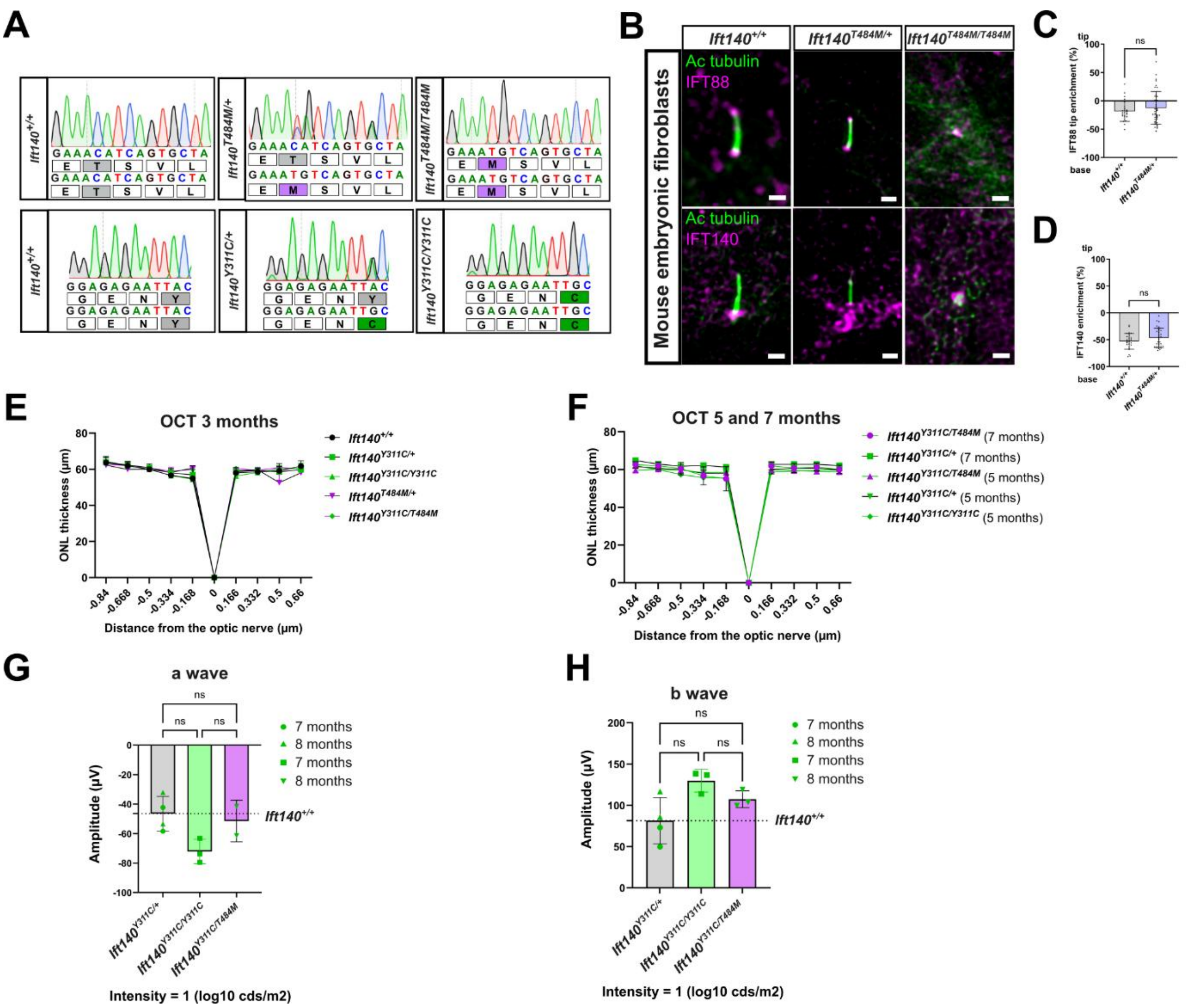


**Supplementary Figure 1. Analysis of *Ift140* knock-in mice harboring the *Ift140* Y311C and T484M missense variants. A)** Sanger sequencing chromatogram showing the change generated in mice to mimic the human T484M and Y311C changes in *Ift140*. **B)** MEFs derived from $Ift140^{+/+}$, $Ift140^{T484M/+}$ and $Ift140^{T484M/T484M}$ mice were stained with the cilium microtubule marker acetylated tubulin (green) and either IFT88 (top panel, magenta) or IFT140 (bottom panel, magenta). Scale bar: 1 µm. The enrichment of immunofluorescence staining at the base of the cilium was quantified for IFT188 (**C**) and IFT140 (**D**). **E and F)** Optical coherence tomography (OCT) measurement of the outer nuclear layer (ONL) thickness from the different *Ift140* genotypes modelled in this study at 3 months (**E**). The number of mice analyzed for each genotype is as follows: +/+ (n=2); Y311C/+ (n=6); Y311C/Y311C (n=3); T484M/+ (n=4); Y311C/T484M (n=2). OCT measurement of the ONL thickness at 5 and 7 months of age (**F**).

The number of mice analyzed for each genotype and time point is as follows: Y311C/+ (5 months; n=6); Y311C/+ (7 months; n=2); Y311C/Y311C (5 months; n=3); Y311C/T484M (5 months; n=3); Y311C/T484M (7 months; n=3). **G-H)** The activity of the retina was measured by ERG, A wave (**G**) showing hyperpolarization of photoreceptors in response to different intensities of light and was plotted as negative values change from baseline. B wave (**H**) showing the response of the inner layer of the retina following phototransduction. The number of mice analyzed for each genotype is as follows: +/+ (n=2); Y311C/+ (7 months, n=2); Y311C/+ (8 months, n=2); Y311C/Y311C (7 months, n=3); Y311C/T484M (8 months, n=3). Kruskal-Wallis test with Dunn’s multiple comparison test was used. Ns = not significant.

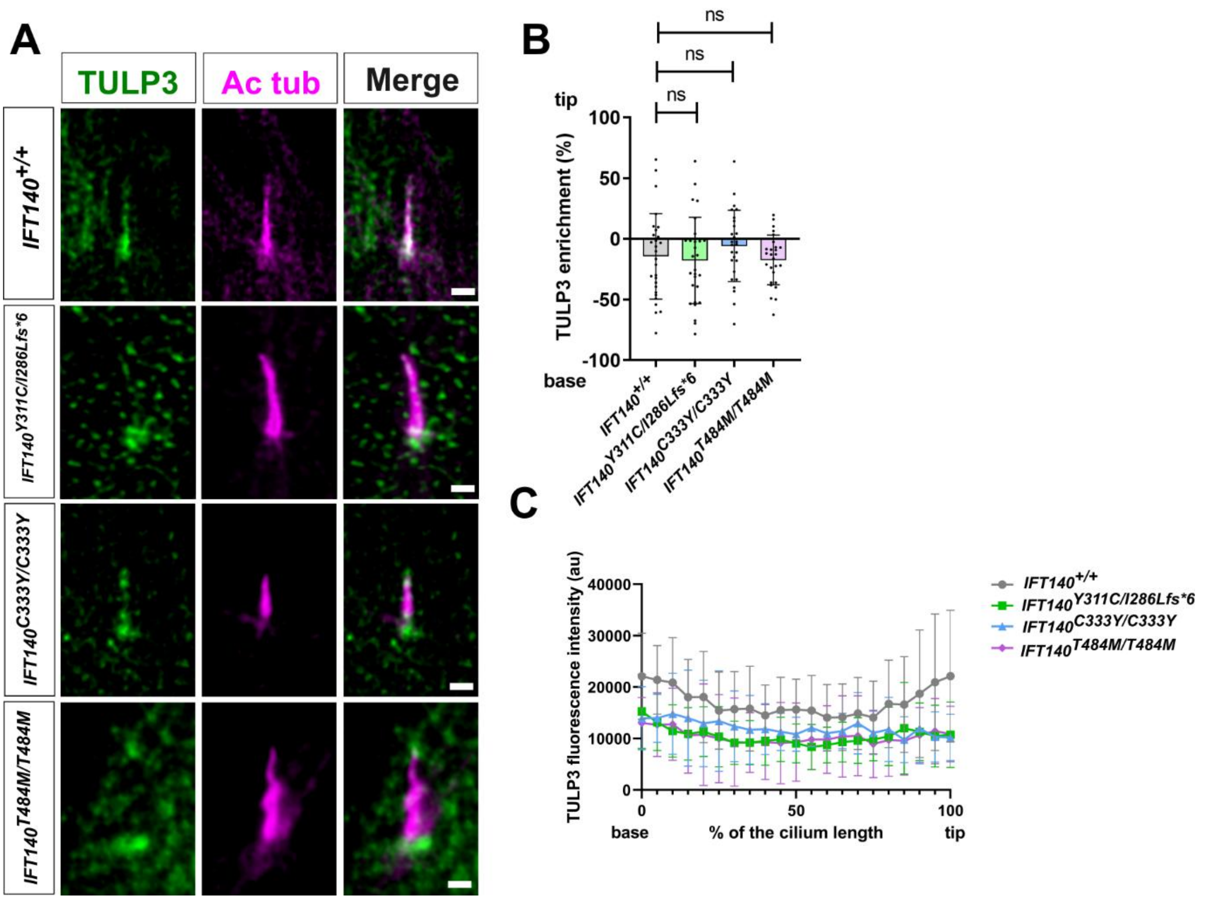


**Supplementary Figure 2. Analysis of TULP3 localization in IFT140 patient fibroblasts. A)** Fibroblasts were stained with TULP3 (green) and acetylated tubulin (magenta) to mark the cilia microtubules. Scale bar: 1 μm. **B)** Percentage of the total TULP3 fluorescence enrichment at the tip is shown for each fibroblast cell line. *IFT140*$^{+/+}$ (n=28 cilia), *IFT140*$^{Y311C/I286Lfs*6}$ (n=28 cilia), *IFT140*$^{C333Y/C333Y}$ (n=27 cilia), *IFT140*$^{T484M/T484M}$ (n=27 cilia). Kruskal-Wallis test with Dunn's multiple comparison test was used. Ns = not significant. **C)** Fluorescence intensity measurements of TULP3 along the cilium of fibroblasts: IFT140$^{+/+}$ (n=28), Y311C (n=30), C333Y (n=22), T484M (n=27).

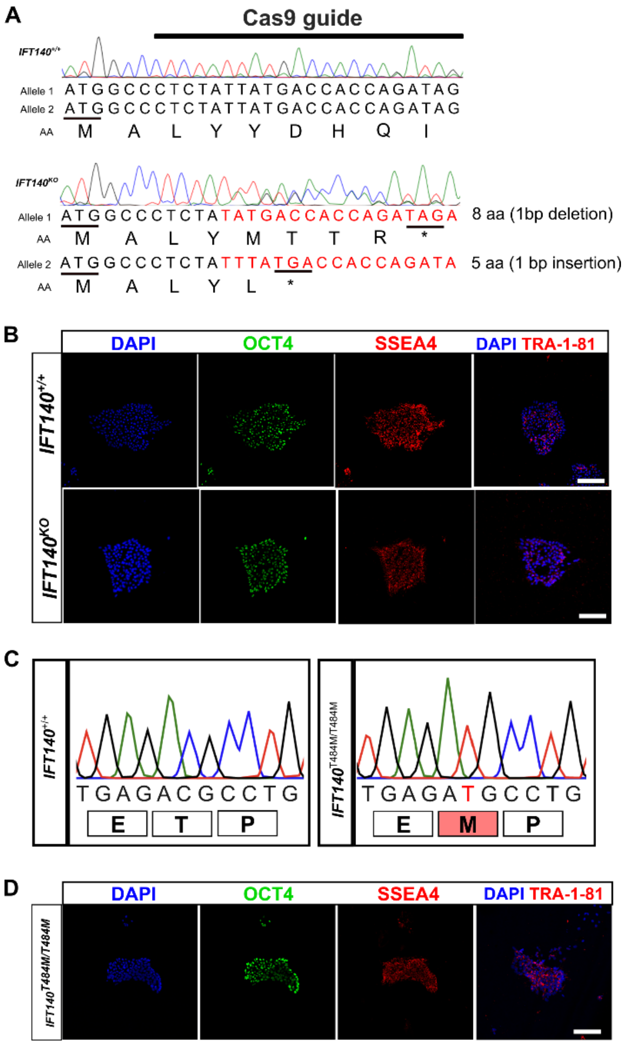


**Supplementary Figure 3. Production and characterization of iPSC lines. A)** Clones were selected following simultaneous CRISPR/Cas9 gene editing and reprogramming on human dermal fibroblasts with gRNA targeting exon 1 of *IFT140*. Sequence traces showing *IFT140* exon 1 in control and *IFT140*$^{KO}$ iPSCs clones. The *IFT140*$^{KO}$ iPSCs clone presents one allele with a 1-bp deletion (MALYMTTR*) and a 1-bp insertion (MALYL*) on the second allele. **C)** Sanger sequence trace of *IFT140*$^{T484M/T484M}$ iPSC clones. **B-D)** Characterization of the IFT140 iPSC clones. **B)** *IFT140*$^{KO}$ and **D)** *IFT140*$^{T484M/T484M}$ iPSC were stained for the pluripotency markers OCT4, SSEA4 and TRA-1-80. DAPI (blue) stains the nucleus. Scale bars: 100 µm.

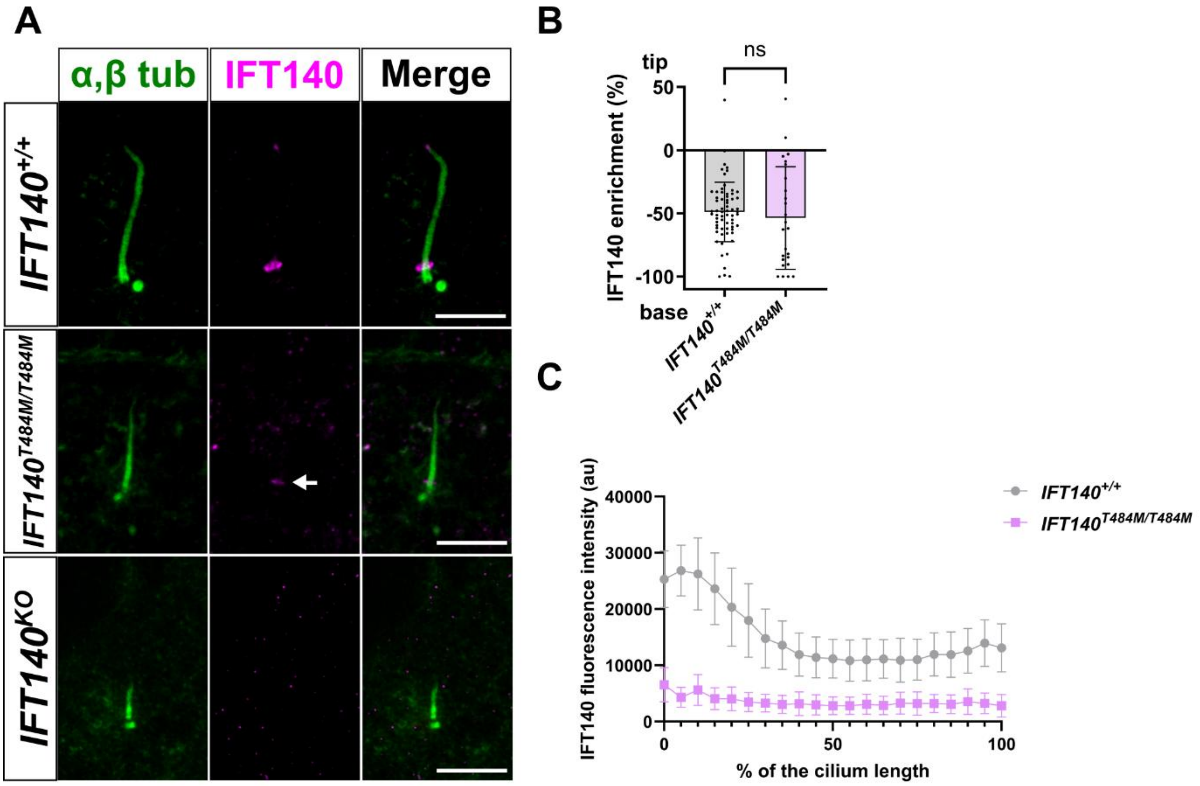


**Supplementary Figure 4. Expansion microscopy analysis of IFT140 localization in iPSC-RPE. A)** U-ExM of iPSC-RPE stained with α and β tubulin (green) and IFT140 (magenta). Scale bar = 10 µm. **B)** Percentage of IFT140 fluorescence intensity enrichment in the cilia base or tip of *IFT140*$^{+/+}$ (n=65 cilia) and *IFT140*$^{T484M/T484M}$ (n=25 cilia) iPSC-RPE. ns = not significant, based on Mann-Whitney test. **C)** Fluorescence intensity measurements of IFT140 along the cilium of iPSC-RPE in *IFT140*$^{+/+}$ (n=65 cilia) and *IFT140*$^{T484M/T484M}$ (n=25 cilia).

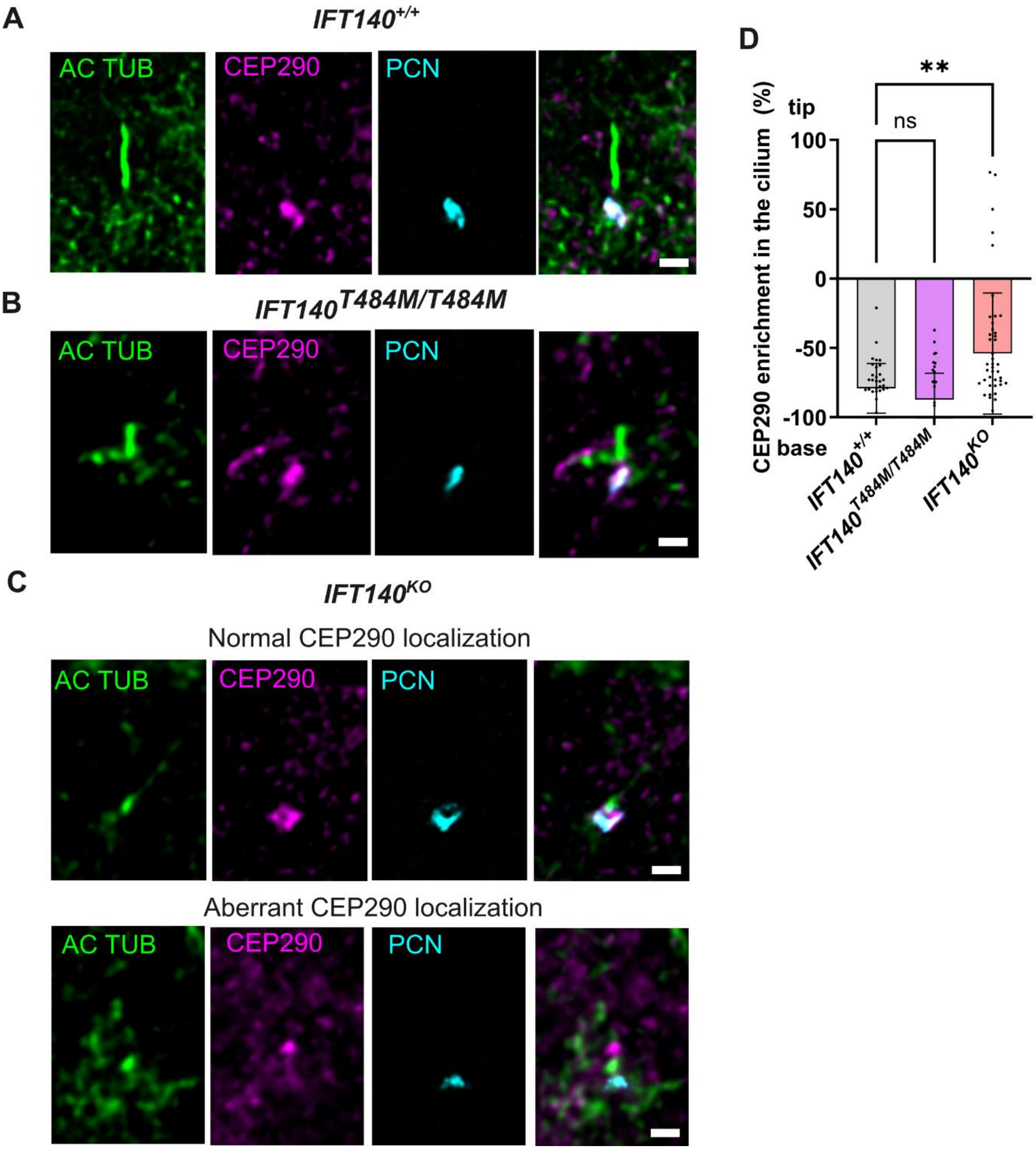


**Supplementary Figure 5. Altered CEP290 localization in *IFT140$^{KO}$* iPSC-RPE.** iPSCs were differentiated to RPE cells and stained with acetylated tubulin (Ac tubulin, green), CEP290 (magenta) and pericentrin (PCN, cyan) **A)** *IFT140$^{+/+}$* **B)** *IFT140$^{T484M/T484M}$* and **C)** *IFT140$^{KO}$*. **C)** shows an example of an *IFT140$^{KO}$* cilium with normal CEP290 localization (top images) and aberrant localization (bottom images). Scale bar: 1 µm. **D)** Percentage of CEP290 fluorescence intensity enrichment in the cilia base or tip of *IFT140$^{+/+}$* (n=39), *IFT140$^{T484M/T484M}$* (n=40 cilia) and *IFT140$^{KO}$* (n=49 cilia) iPSC-RPE. ns = not significant, ** p-value <0.001 based on Kruskal-Wallis test with Dunn's multiple comparison test.

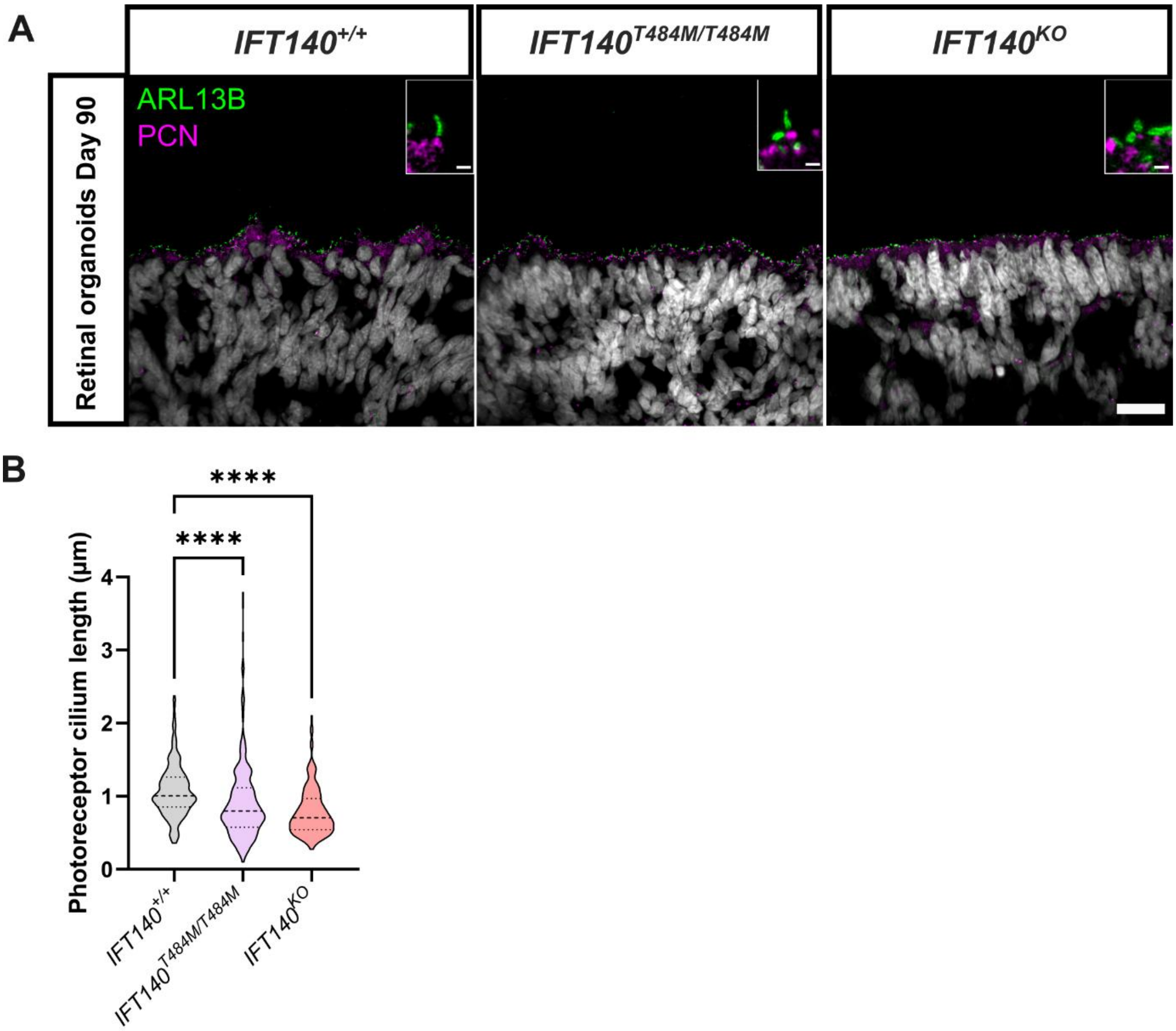


**Supplementary Figure 6. Photoreceptor cilium length is reduced in IFT140 iPSC-ROs. A)** Retinal organoids were differentiated from iPSCs for 90 days and stained with the cilium marker ARL13B (green), the basal body marker PCN (magenta) and DAPI (grey). Scale bar: 20 µm. The insets show a magnification of the connecting cilium region. Scale bar: 1 µm. **B)** Photoreceptor cilium length was measured manually using the length of ARL13B. Data from *IFT140*$^{+/+}$ (n=132 cilia, n=3 ROs), *IFT140*$^{T484M/T484M}$ (n=278 cilia, n=4 ROs) and *IFT140*$^{KO}$ (n=148 cilia, n=3 ROs). **** p-value <0.0001 based on Kruskal-Wallis test with Dunn's multiple comparison test.

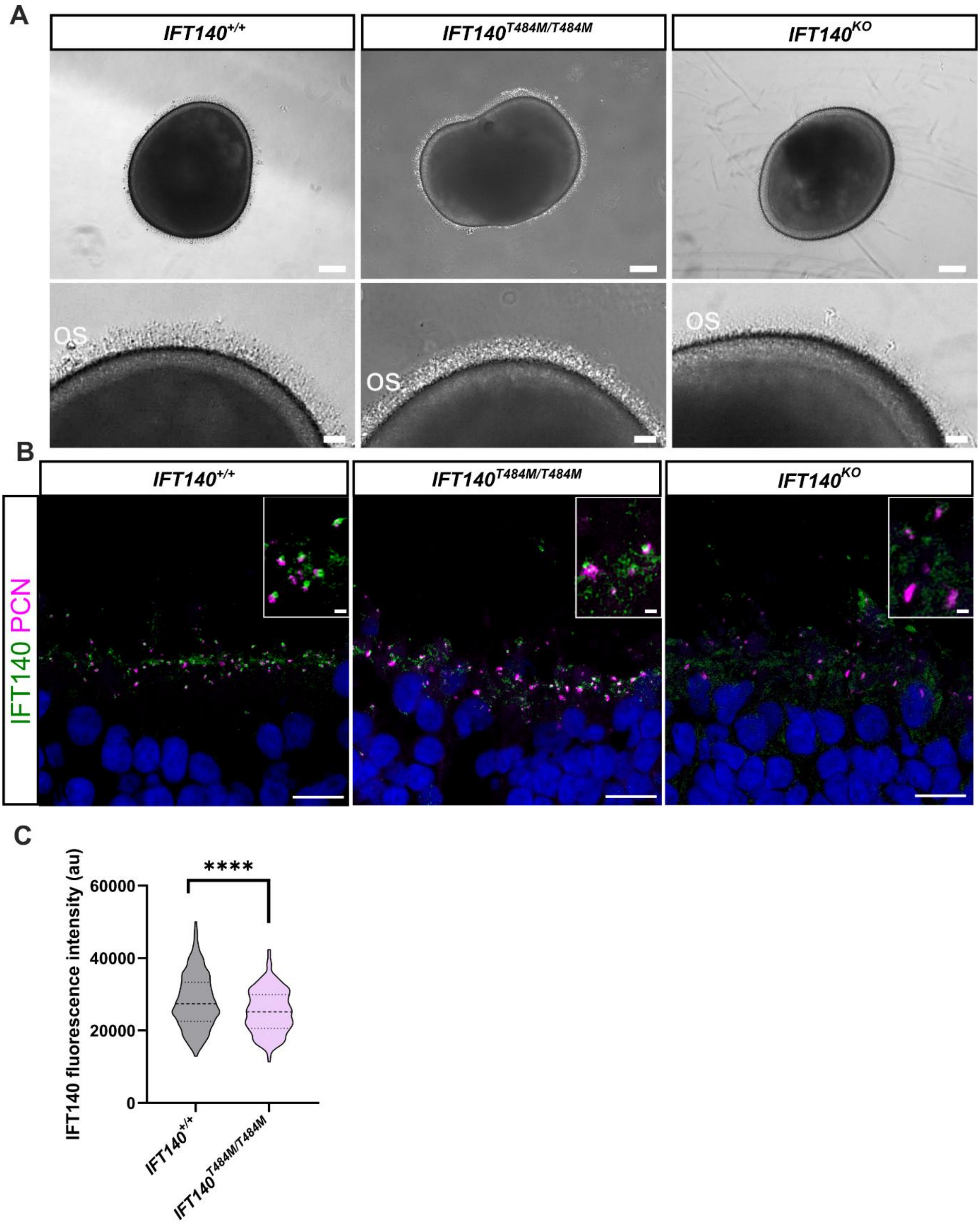


**Supplementary Figure 7. IFT140 immunofluorescence is reduced in iPSC-ROs.** iPSCs were differentiated to retinal organoids for 200 days, embedded and cryosectioned. **A)** Brightfield images of the retinal organoids at day 200, showing the presence of outer segments (OS) in all lines. Scale bar: 200 µm for upper panels, 50 µm for lower panels. **B)** Cryosections were stained with an IFT140 antibody (green) and PCN (magenta). Scale bar: 10 µm for lower magnification images, 1 µm for insets. **C)** IFT140 fluorescence intensity in the cilium was measured for *IFT140*$^{+/+}$ (n=326 cilia, n=5 ROs) and *IFT140*$^{T484M/T484M}$ (n=186 cilia, n=4 ROs) iPSC-ROs. **** p-value <0.0001, based on Mann-Whitney test.

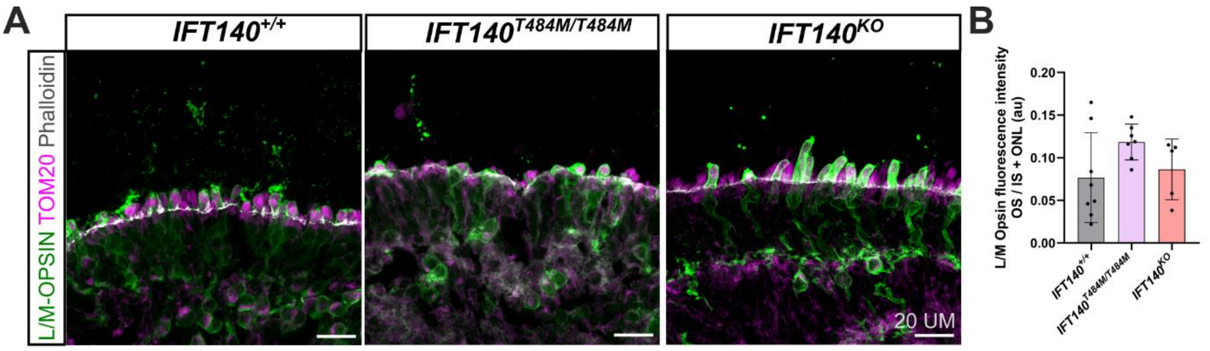


**Supplementary Figure 8. L/M opsin localization is not altered in IFT140 iPSC-ROs.** Retinal organoids from the *IFT140*$^{KO}$ line, isogenic controls, and the *IFT140*$^{T484M/T484M}$ patient line were collected at day 200 post differentiation, embedded and cryosectioned. Data from n=2 independent differentiations. **A)** Staining of L/M opsin (green) and TOM20 (magenta) marking the inner segments and phalloidin (white) marking the outer limiting membrane. Scale bar: 20 µm. **B)** The fluorescence intensity of L/M opsin was measured in the outer segments (OS) and the inner segment and outer nuclear layer region (IS+ONL), and the ratio OS/IS+ONL was obtained. The analysis consisted of: *IFT140*$^{+/+}$ (n=8 images from n=4 ROs); *IFT140*$^{T484M/T484M}$ (n=7 images from n=5 ROs); *IFT140*$^{KO}$ (n=5 images from n=3 ROs).

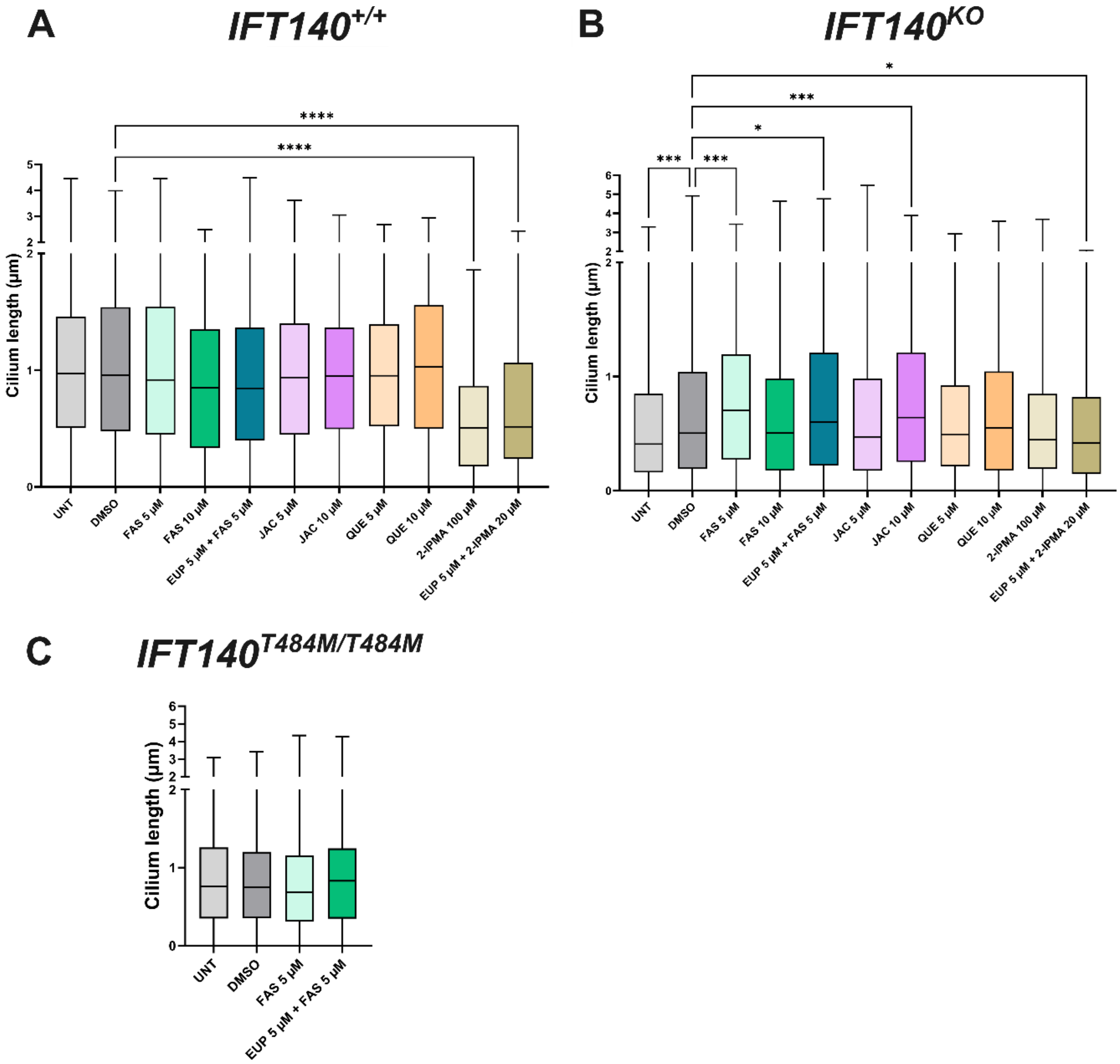


**Supplementary Figure 9. Small molecule treatment of iPSCs-RPE.** Cilium length in **A)** control (*IFT140*$^{+/+}$) **B)** *IFT140*$^{KO}$ and **C)** *IFT140*$^{T484M/T484M}$ iPSC-RPE following treatment with the following compounds: UNT (untreated), DMSO (vehicle), FAS (fasudil), JAC (jaceosidin), QUE (quercetin) or 2-isopropylmalic acid (2-IPMA). Cells were treated for 24 hours and stained using ARL13B and PCN. Cilia length was measured using CiliaQ. At least 200 cilia were measured for each condition; ≥2 independent differentiations; and ≥3 independent experiments were performed for each condition. Significant differences between the treatment and the DMSO condition are shown as $p<0.05$ (*), $p<0.01$ (**), $p<0.001$ (***), $p<0.0001$ (****), based on Kruskal-Wallis test with Dunn's multiple comparison test.

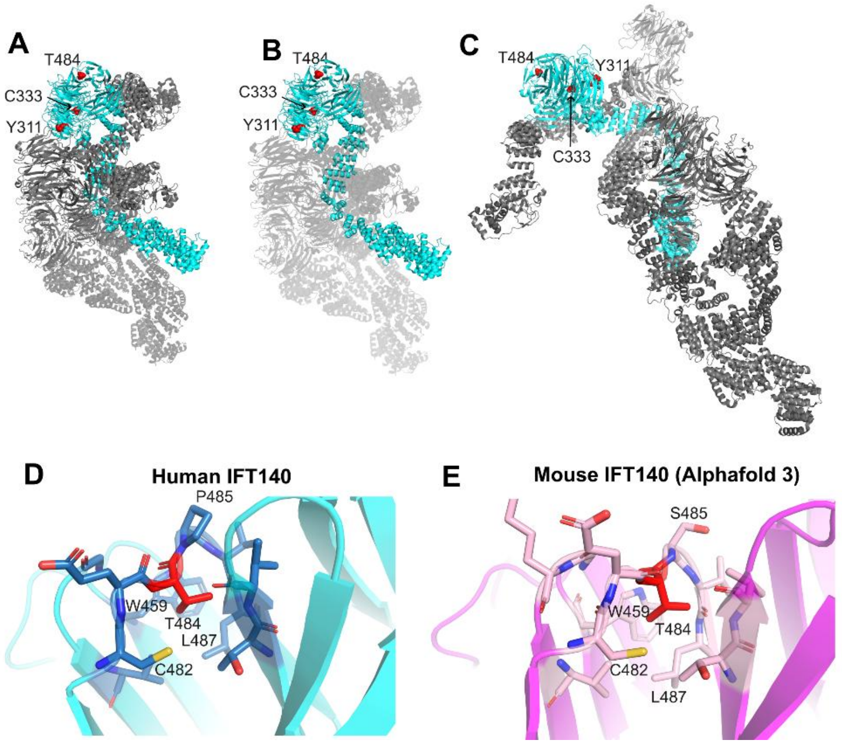


**Supplementary Figure 10. Localization of IFT140 patient variants within the IFT-A complex.** Cryo-EM structure of human IFT-A (PDB 8BBG). IFT140 is shown in cyan; all other subunits (IFT144, IFT122, IFT139, IFT121, IFT43) in grey. The three missense variant positions from this study (Y311, C333, T484) are shown as red spheres. All three residues are buried within the N-terminal tandem WD40 β-propeller domain of IFT140 and are not located at inter-subunit interfaces. **A)** Full complex, all subunits opaque. **B)** IFT140 opaque, other subunits semi-transparent (60% transparency) to emphasize IFT140 within the complex. **C)** Different orientation of the complex to show that the three variants studied are not in interaction interface. **D-E)** Structural environment of T484 in human and mouse IFT140. Close-up view of T484 and surrounding residues shown as sticks. T484 (red) is located on a β-strand in WD40 propeller 2. **D)** Human IFT140 (cryo-EM, PDB 8BBG chain B, cyan). The label P485 indicates the adjacent proline. **E)** Mouse IFT140 (AlphaFold3 model, pink/magenta). The label S485 indicates the adjacent serine, which is the key human–mouse sequence difference at this position. The T484M substitution (threonine → methionine) introduces a larger, hydrophobic side chain into a buried position, with a predicted destabilisation of +1.23 kcal/mol (human) and +1.48 kcal/mol (mouse) by ThermoMPNN, the largest effect among the three missense variants.

### Supplementary Tables

**Supplementary Table 1. Predicted thermodynamic effect of IFT140 missense variants.** ΔΔG values predicted by ThermoMPNN (kcal/mol; positive = destabilising). Values computed on the cryo-EM structure (PDB 8BBG chain B); mouse values on the AlphaFold3 top-ranked model. All three variants are mildly destabilizing, with T484M showing the strongest effect in both species.

| IFT140 variant | Hs ΔΔG (kcal/mol) | Mm ΔΔG (kcal/mol) |
|---|---|---|
| Y311C | +0.80 | +0.74 |
| C333Y | +0.69 | +0.52 |
| T484M | +1.23 | +1.48 |

**Supplementary Table S2. Donor oligonucleotide sequences used for the generation of the Ift140 knock-in mouse lines.** Mutant sequences are underlined; synonymous mutation sequences are in bold.

| Mouse protein change | Donor oligonucleotide sequence |
|---|---|
| p.Y311C | CTTCACAGCACTTTCCCTCTTTATATTGCAGGTTCTGGGACTTAGAACGGGGAGAGAATTGC**ATCTTA**AGT**CTCCAA**GAGAAGTTTGGCTTTGAAAAGGGAGAGAGTATAAACTGTGTGTGTTTCTGTAAAGCCAAA |
| p.T484M | CACCACGCTGCTGCTGTTTCAATACTCAATCTTTCATTTCAGGGACATTCCTGTGTGAAATGTCA**GTA**CTAGCCATGCAT**GAA**GAAAGCATTTACACCGTGGAGCCAAACCGACTCCAAGTCCGGACCTGGCAGGTAAGTGAC |

**Supplementary Table S3. List of genotyping primers used in this study.**

| Gene (human) | Primer | |
|---|---|---|
| | Forward | Reverse |
| *IFT140 exon 13* (for T484M variant) | TAGTGCACCCCTATGAAGCT | AACTTACCTGCCAGGTTCGA |

| | | |
|---|---|---|
| *IFT140 exon 9* (for Y311C and C333Y variants) | GGCTGCAATTCCTGTGTTC | AACCAACCACAAGCATTGT |

| **Gene (mouse)** | **Primer** | |
|---|---|---|
| | **Forward** | **Reverse** |
| *Ift140 exon 12* (for T484M variant) | CCGTCGTTCCCTGTGTTCTGAG | TACCCCGGGGGCCGTATTTT |
| *Ift140 exon 8* (for Y311C variant) | ATATGTAGCTGGGACTCCAAT | TTAAAACATAGACCAGGCTG |
| *rd8* | GGTGACCAATCTGTTGACAATCC | GCCCCATTTGCACACTGATGAC |
